# CD81^+^ senescent-like fibroblasts exaggerate inflammation and activate neutrophils via C3/C3aR1 axis in periodontitis

**DOI:** 10.1101/2024.02.21.581332

**Authors:** Liangliang Fu, Chenghu Yin, Qin Zhao, Shuling Guo, Wenjun Shao, Ting Xia, Quan Sun, Liangwen Chen, Jinghan Li, Min Wang, Haibin Xia

**Affiliations:** State Key Laboratory of Oral & Maxillofacial Reconstruction and Regeneration, Key Laboratory of Oral Biomedicine Ministry of Education, Hubei Key Laboratory of Stomatology, School & Hospital of Stomatology, Wuhan University, Wuhan 430079, China; (L.F.); (C.Y.)

**Keywords:** Cellular senescence, Periodontal diseases, Human gingival fibroblasts, Senescence associated secretory phenotype, CD81

## Abstract

Periodontitis, a prevalent inflammatory disease worldwide, poses a significant economic burden on society and the country. Previous research has established a connection between cellular senescence and periodontitis. However, the role and mechanism of cell senescence in the progression of periodontitis have not been thoroughly investigated. This study aimed to explore the involvement of cellular senescence in the pathogenesis of periodontitis and determine the underlying mechanisms. Our findings demonstrated that senescent cells accumulated during the progress of periodontitis. Moreover, several scRNA-seq analysis suggested that gingival fibroblasts were the main cell population undergoing cellular senescence during periodontitis, which helps mitigate tissue damage and bone loss. Furthermore, we identified a high expression of CD81 in the senescent gingival fibroblast population. These cells were found to actively contribute to inflammation through their potent pro-inflammatory metabolic activity and secretion of SASP-related factors. Additionally, they recruited neutrophils via the C3/C3aR1 pathway, indirectly sustaining the inflammatory response. Senolytics via Navitoclax successfully alleviated inflammation and bone loss in periodontitis and administration of metformin could alleviate alleviated inflammation and bone loss in periodontitis through inhibiting cellular senescence. These results provide valuable insights into the cellular and molecular basis of periodontitis-induced tissue damage, highlighting the significance of fibroblast senescence. In conclusion, our study sheds light on the relationship between CD81 and cellular senescence, suggesting its potential as a therapeutic target for periodontitis.

## 1. Introduction

Periodontitis is an inflammatory disease of irreversible progressive tissue damage, alveolar bone loss and destruction of tooth supporting tissues, and is caused by microbial infections that eventually lead to tooth loosening and eventual tooth loss (Wolff, Dahlén, & Aeppli, 1994). Periodontitis affects 11.2% of the global population and more than 40% of people over the age of 30, posing a major burden on public health (Eke, Borgnakke, & Genco, 2020; Sanz et al., 2020). Clinical studies have shown that the prevalence and severity of periodontitis increase with age, and moderate loss of alveolar bone and periodontal attachment is common in older adults (Huttner, Machado, de Oliveira, Antunes, & Hebling, 2009).

Cell senescence is a stress response characterized by irreversible proliferation arrest, resistance to apoptosis, and secretion of a range of inflammatory cytokines, growth factors, and proteases, known as senescence-associated secretory phenotypes (SASP) (Coppé, Desprez, Krtolica, & Campisi, 2010; Rodier et al., 2009). Cellular senescence is considered necessary for tissue homeostasis as it aims to eliminate unnecessary damage and promote tissue repair through immune-mediated mechanisms, and even prevent the occurrence of tumors (Campisi, 2013; Ohtani & Hara, 2013). However, the specific environment of the gingival sulcus leads to persistent plaque in periodontal tissue, resulting in oxidative DNA damage as collateral damage of chronic bacterial infection (Aquino-Martinez et al., 2020). Repeated exposure to lipopolysaccharide (LPS) derived from *Porphyromonas gingivalis* (Pg), a key pathogen of periodontitis, can accelerate cellular senescence driven by DNA damage (Aquino-Martinez et al., 2021). Furthermore, recent evidence suggests that bacteria can also induce senescence of healthy cells in an active oxygen-dependent manner by causing inflammation and excessive neutrophil activity (Guo et al., 2024; Lagnado et al., 2021). The aggravation or persistence of these stimulating factors can lead to abnormal accumulation of senescent cells and directly affect periodontal tissue function. Therefore, chronic bacterial infections can cause cell senescence through both direct and indirect mechanisms.

Senescent cells have been found to contribute to bacteria-induced inflammation, with the activation of SASPs playing a crucial role in the release of various pro-inflammatory factors, including interleukin (IL)-1α, IL-6, and IL-8. Elevated levels of these inflammatory factors have been associated with periodontal damage and loss of alveolar bone (Aquino-Martinez et al., 2020; Yu et al., 2024). However, the specific mechanism by which senescent cells contribute to the development of periodontitis remains unclear. In the immune response to periodontitis, dendritic cells (DCs) infected by Pg activate related SASPs, such as IL-1β, IL-6, and IL-8, which ultimately accelerate the progression of periodontitis (El-Awady, Elashiry, Morandini, Meghil, & Cutler, 2022). Additionally, the aging of T lymphocytes, which are crucial for adaptive immunity, leads to a significant alteration in their immunosuppressive ability in Th17/Treg subsets. This alteration ultimately results in the loss of tooth support and alveolar bone (González-Osuna et al., 2022). However, the role and mechanism of cellular senescence in the progression of periodontitis have not been thoroughly investigated.

The breakthrough technology of single-cell RNA sequencing has made it easier to analyze gene expression at the cellular level and identify key cell subpopulations (Zhang et al., 2021). In this study, we utilized bulk-RNA seq, clinical periodontal samples, and a mice ligature-induced periodontitis model to demonstrate that cellular senescence levels increase with periodontitis progression. Through sc-RNA seq, in vitro, and in vivo experiments, we observed significant cellular senescence in gingival fibroblasts. Additionally, we identified a unique subgroup of gingival fibroblasts with high expression of CD81, which exhibited senescence characteristics such as ROS accumulation and enrichment of senescence genes. We propose that this subgroup of fibroblasts can directly promote the progression of periodontitis by secreting SASP-related factors, such as IL-6, and indirectly amplify inflammation by recruiting neutrophils through the complement pathway, specifically C3. We also found that targeting cellular senescence with senolytic drug or metformin can reduce inflammation and delay alveolar bone resorption in periodontitis.

## 2. Results

### 2.1 Cellular senescence characteristics in periodontitis

Cellular senescence is a manifestation of aging at the cellular level. Although accumulation of senescent cells is normal in aged tissues, persistent bacterial infection and chronic inflammation promote the early onset of senescence by ROS activation and DNA damage (Aquino-Martinez, 2023). In clinical gingival specimens from periodontally healthy individuals of similar age and those diagnosed with periodontitis, we found that the senescence biomarker senescence-associated β-galactosidase (SA-β-gal) was scarcely expressed in the gingiva of young healthy individuals. However, in gingival samples from patients with periodontitis, a notable increase in SA-β-gal-positive cells was observed, primarily localized in the lamina propria of gingival connective tissue (Figure 1A). Additionally, IHC staining analysis revealed that other senescent biomarkers, such as cell cycle inhibitory proteins p16 and p21, and senescence-associated heterochromatin foci (SAHFs) like H3K9me3, were significantly upregulated in human periodontitis gingival tissues as well (Figure 1B).

**Figure 1.**
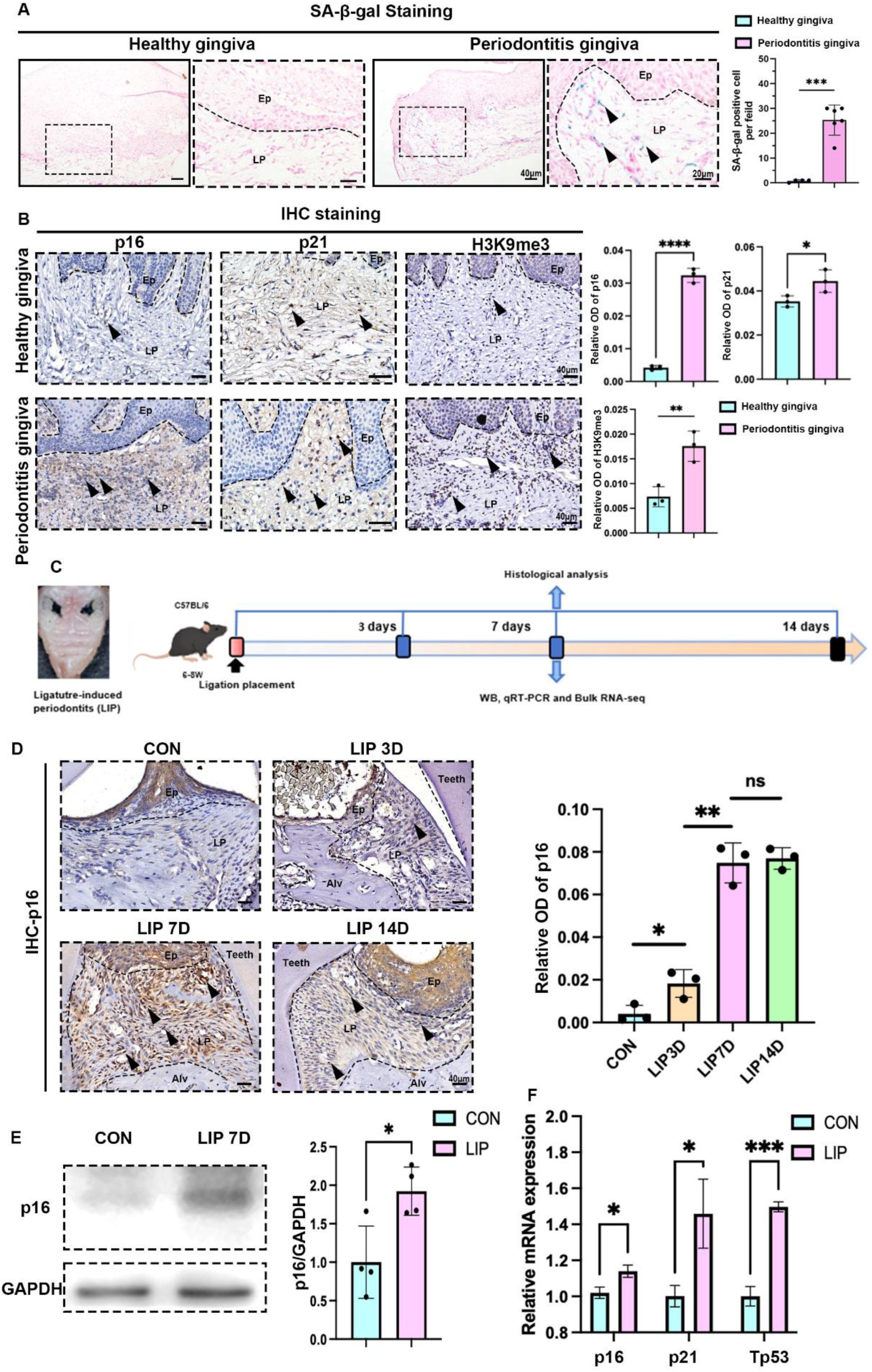
Characteristics of cellular senescence along with periodontitis progression. (**A**) Representative image of and semi-quantification of SA-β-gal staining in healthy (n=4 field) and periodontitis (n=6 field) patient gingiva, scale bar=40μm or 20μm. (**B**) Representative images of IHC staining and semi-quantification of p16, p21 and H3K9me3 in healthy and periodontitis patient gingiva (n=3 field), positive cells were indicated by black arrow, scale bar=40μm. (**C**) Analysis strategy of ligature-induced periodontitis (LIP) mouse model. (**D**) Representative image of IHC staining and semi-quantification of p16 in mouse gingiva of health and LIP post 3, 7 and 14 days (n=3 field), scale bar=40μm. (**E**) Western blot images and semi-quantification of p16 protein levels in control (CON) and LIP post 7 days (LIP 7D) mouse gingiva (n=4 independent experiments). (**F**) qrt-PCR analysis of p16, p21and Tp53 in control (CON) and LIP 7D mouse gingiva (n=3 independent experiments). Ep: Epithelium; LP: Lamina propria; Alv: Alveolar bone; Teeth. Data are expressed as mean ± SD. *p<0.05, **p<0.01, ***p<0.001. ****p<0.0001.

The clinical samples from periodontitis patients were often derived from older individuals, because periodontitis incidence obviously increases with age (Eke et al., 2020). To avoid confounding factors like age potentially affecting the experimental results, we also examined the levels of cellular senescence in the Ligature-induced periodontitis (LIP) mouse model (Figure 1C). IHC staining results indicated that the protein expression level of p16 among gingiva was significantly upregulated following ligation, peaking at day 7 post-ligation (Figure 1D). And then, gingiva at day 7 post-ligation and healthy gingiva as control were collected for protein and gene analysis. Western blotting analysis showed that the protein levels of p16 in LIP 7D group was about 2 times of that in control group (Figure 1E). And the transcription of p16, p21, and Tp53 in gingival tissues were higher at day 7 post-ligation than those in control (Figure 1F). Furthermore, bulk RNA-sequencing was performed on gingival tissues from LIP 7D and healthy mice (Figure 1-figure supplement 1A), identifying 458 upregulated and 358 downregulated genes. Notably, among the upregulated genes, 19 senescence-associated genes were detected, including C3, Il6 and so on (Figure 1-figure supplement 1B). We also observed a significant upregulation of several SASPs genes such as Icam1, Mmp3, Nos3, Igfbp7, Igfbp4, Mmp14, Timp1, Ngf, Il6, Areg, and Vegfa in the LIP group (Figure 1-figure supplement 2A). The Gene Set Enrichment Analysis (GSEA) enrichment analysis based on our sequencing data revealed the upregulation of the cellular senescence pathway in LIP mice Figure 1-figure supplement 1C). Moreover, a significant reduction in oxidative phosphorylation and the tricarboxylic acid (TCA) cycle was observed in the LIP group (Figure 1-figure supplement 2B, C).

Gene Ontology Biological Process (GO BP) analysis of differentially expressed genes further demonstrated mitochondrial respiratory and electron transport dysfunction, as well as impaired oxidative phosphorylation in the gingiva of LIP mice, suggesting that mitochondrial dysfunction might contribute to cell senescence in periodontitis (Figure 1-figure supplement 1D). Meanwhile, upregulation of the cellular senescence pathway and a series of inflammatory-related pathways, including complement activation and response to lipopolysaccharide, were also enriched in the LIP group (Figure 1-figure supplement 1D). Besides that, the PI3K-AKT, MAPK and NF-κB signaling pathways were also activated in LIP group (Figure 1-figure supplement 2D, E and F), which were closely associated with the onset of cellular senescence and the secretion of SASP factors (Raynard et al., 2022; Sayegh et al., 2024; Tang et al., 2023). Collectively, these findings suggested that senescent cells gradually accumulated and senescence-related signaling pathways were activated during the progression of periodontitis.

### 2.2 Gingival fibroblasts were the main cell type responsible for cellular senescence in periodontitis

To identify which cell types in periodontitis tissue are enriched for senescence, we re-analyzed public sc-RNA seq data of healthy and periodontitis human gingiva (Williams et al., 2021). This data from 8 healthy and 13 periodontitis-affected gingival samples were analyzed, clustering the cells into 15 distinct groups (Figure 2-figure supplement 1A). These clusters were classified into fibroblasts, immune cells, epithelial cells, endothelial cells, and other cell types based on specific markers (Figure 2A, Figure 2-figure supplement 1B).

**Figure 2.**
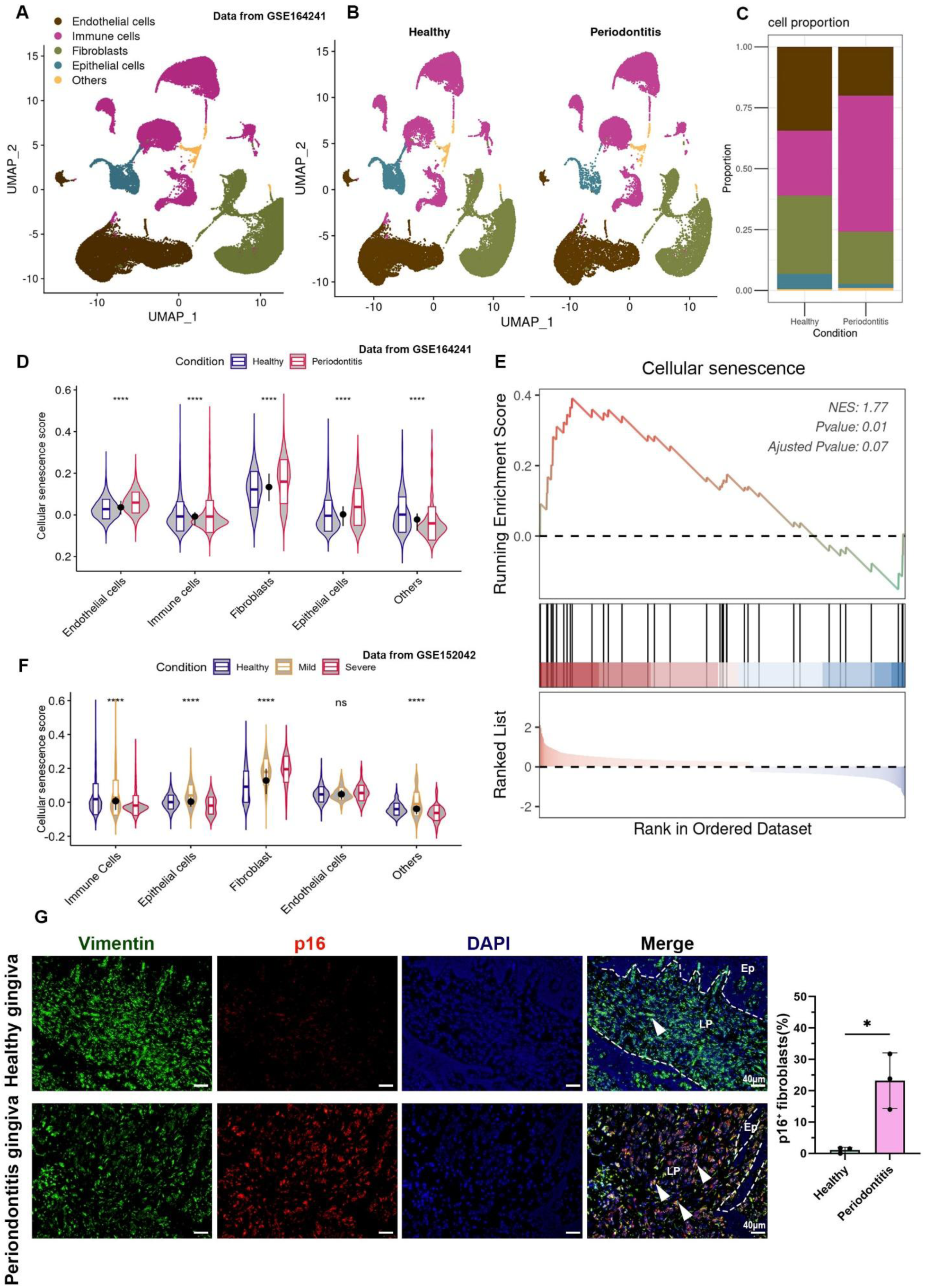
Cellular Senescence of gingival fibroblasts in periodontitis. **(A and B)** UMAP diagram and single-cell annotation of cells clusters for the healthy and periodontitis patient gingiva from public dataset GSE164241. (**C**) Histogram of gingival tissue cell ratio in healthy and periodontitis patients. (**D**) The violin plot showing cellular senescence score of cell groups in healthy and periodontitis gingiva. (**E**) GSEA enrichment analysis of cellular senescence pathway in fibroblasts among periodontitis compared to those in healthy gingiva. (**F**) The violin plot showing cellular senescence score in subgroups in gingiva of healthy, mild and severe periodontitis patient from public dataset GSE152042. (**G**) Immunofluorescence staining and semi-quantification of p16 positive fibroblasts in healthy and periodontitis patient gingiva. p16 (red), Vimentin (green), and nuclei (blue), Ep: Epithelium; LP: Lamina propria. White arrow indicates double positive cells, scale bar=40μm, n=3.

In periodontitis samples, there was a notable shift in cellular composition: immune cells increased while structural cells, such as fibroblasts, decreased (Figure 2B, C). Cellular senescence gene score analysis across different cell types revealed that fibroblasts in particular showed significant upregulation of senescence scores in periodontitis, indicating that they had the highest overall levels of senescence (Figure 2D). GSEA of differentially expressed genes between healthy and periodontitis fibroblasts further confirmed the activation of senescence pathways in periodontitis (Figure 2E). To further verify fibroblast senescence in periodontitis, we analyzed another data from GSE152042, which included samples from 2 healthy, 1 mild, and 1 severe periodontitis gingiva (Caetano et al., 2021). The results showed a decline in fibroblast proportion along with increasing disease severity (Figure 2-figure supplement 1C and D) and a corresponding increase in cellular senescence score (Figure 2F). Immunofluorescence staining on clinical sample confirmed that the proportion of P16^+^ senescent fibroblasts in periodontitis rose to approximately 25%, compared to very few in healthy gingiva (Figure 2G). *In vitro,* healthy primary gingival fibroblasts (HGFs) stimulated with different concentrations of *Porphyromonas gingivalis* lipopolysaccharide (Pg-LPS) showed a dose-dependent increase in SA-β-gal positive fibroblasts (Figure 2-figure supplement 2A and B). These findings suggest that gingival fibroblasts undergo significant senescence, potentially induced by Pg-LPS, during the progression of periodontitis.

### 2.3 CD81^+^ fibroblasts were identified as the major fibroblast subpopulation undergoing senescence

To examine the changes in gingival fibroblast subpopulations during periodontitis, we analyzed gingival fibroblasts from dataset GSE164241 (Williams et al., 2021), and identified 7 distinct fibroblasts subpopulations (Figure 3A). The cell proportion bar chart revealed a significant increase in subpopulations 1 and 3 in periodontitis compared to healthy controls (Figure 3B). We then applied a cellular senescence gene set (Saul et al., 2022), to score these subpopulations and found that subpopulation 1 exhibited the highest average expression levels, with a marked increase in periodontitis (Figure 3C). GO enrichment analysis of the differentially expressed genes further confirmed that subpopulation 1 displayed upregulated aging characteristics Figure 3D), indicating that this subpopulation is primarily responsible for fibroblast senescence. Among the top 20 marker genes for subpopulation 1, CD81, a transmembrane protein, emerged as a potential biomarker for this senescent subpopulation (Figure 3E). A density heatmap demonstrated that CD81 was predominantly enriched in subpopulation 1 (Figure 3F). The remaining subgroups were classified as EmFB (extracellular matrix-associated fibroblasts), P-EmB (pre-extracellular matrix-associated fibroblasts), MyFB (myofibroblasts), P-MyFB (pre-myofibroblasts), VFB (vascular-associated fibroblasts), and ImFB (immune-associated fibroblasts), based on GO analysis (Figure 3F). Immunofluorescence staining further showed that the proportion of CD81^+^ fibroblasts in periodontitis increased to approximately 50%, compared to very few in healthy samples (Figure 3G). Thus, CD81^+^ fibroblasts might represent a core senescent fibroblast population in human periodontitis.

**Figure 3.**
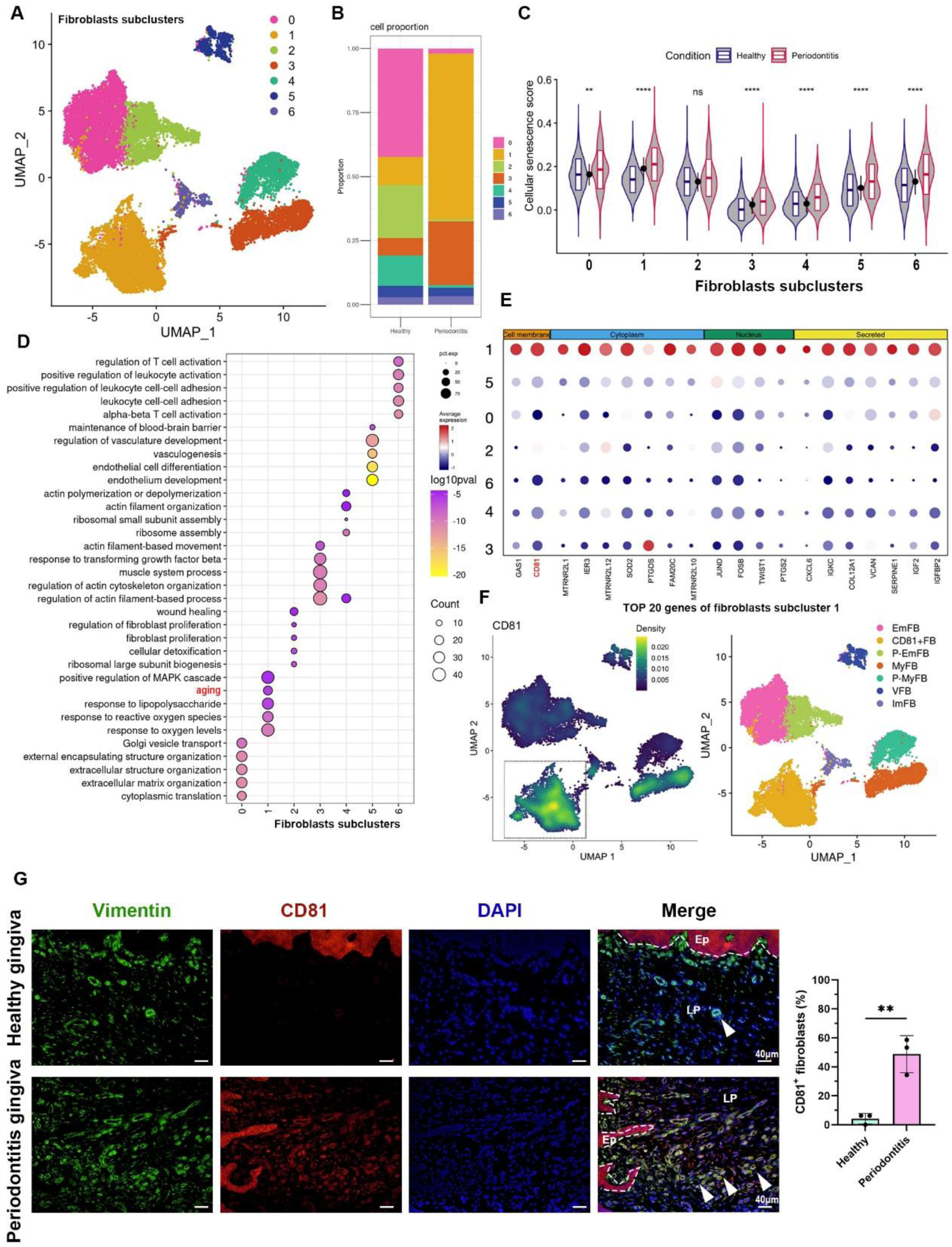
CD81 is identified as the potential marker of senescent gingival fibroblast. (**A**) UMAP diagram illustrated the cell subclusters of fibroblasts from public dataset GSE164241. (**B**) Histogram of fibroblasts subclusters ratio in healthy and periodontitis gingiva respectively. (**C**) The violin plot showing cellular senescence score of each fibroblast subcluster in healthy and periodontitis gingiva. (**D**) GO enrichment analysis of each fibroblast subcluster. Fibroblast subcluster 1 shows enrichment of aging process highlighted by red. (**E**) Cellular localization of the top 20 marker molecules in fibroblasts subcluster 1. CD81 protein, located at cell membrane, was highlighted by red. (**F**) Density map of CD81 expression among fibroblast subcluster and re-annotation of fibroblast subcluster according to GO analysis. (**G**) Immunofluorescence staining and semi-quantification of CD81 positive fibroblasts in healthy and periodontitis patient gingiva. VIM (green), CD81 (red) and nuclei (blue), Ep: Epithelium; LP: Lamina propria. White arrow indicates double positive cells, scale bar=40μm, n=3. Data are expressed as mean ± SD. * P <= 0.05, **P<= 0.01, ***P<= 0.001, ****P<= 0.0001.

### 2.4 CD81^+^ fibroblasts were terminally differentiational cell with high SASP expression

To investigate the role of fibroblasts in periodontitis-related inflammation, we analyzed the expression of SASP-related genes in each fibroblast group. CD81^+^ fibroblasts exhibited elevated levels of SASP-related genes, including IL-6, CXCL-5, CXCL-6, MMP1, and MMP3 (Figure 4A). Additionally, we examined the metabolic activity of each subgroup, focusing on lipid metabolism. Pathways related to fatty acid biosynthesis, arachidonic acid metabolism, and steroid biosynthesis were significantly upregulated in CD81^+^ fibroblasts (Figure 4-figure supplement 1A), suggesting that lipid metabolism might play a role in cellular senescence of the gingival fibroblasts. Arachidonic acid, in particular, could be converted into prostaglandins and leukotrienes via cyclooxygenases (COXs) and lipoxygenases, contributing to the inflammatory response (Figure 4-figure supplement 1B) (Wang et al., 2021). We further observed a higher gene expression of PTGS1 (encoding COX1 protein) and PTGS2 (encoding COX2 protein) in CD81^+^ fibroblasts compared to other fibroblasts subpopulations (Figure 4-figure supplement 1C). Pseudotime analysis of fibroblast differentiation trajectories revealed that CD81^+^ fibroblasts predominantly clustered at the end of the trajectory, indicating limited differentiation potential (Figure 4B, C). Functional enrichment analysis of genes showing gradual increases during differentiation highlighted pathways related to inflammatory activation and aging characteristics (Figure 4D). Several SASP genes, including CXCL1, CXCL6, IL6, MMP1, SERPINE1, EGFR, FGF2, FNDC1, IGFBP4, LAMB1, and TIMP1, also exhibited increased expression during differentiation (Figure 4E). Overall, our bioinformatics analysis demonstrated that CD81^+^ fibroblasts exhibited differentiation arrest and heightened expression of SASP factors, further implicating them in the inflammatory and senescent processes of periodontitis.

**Figure 4.**
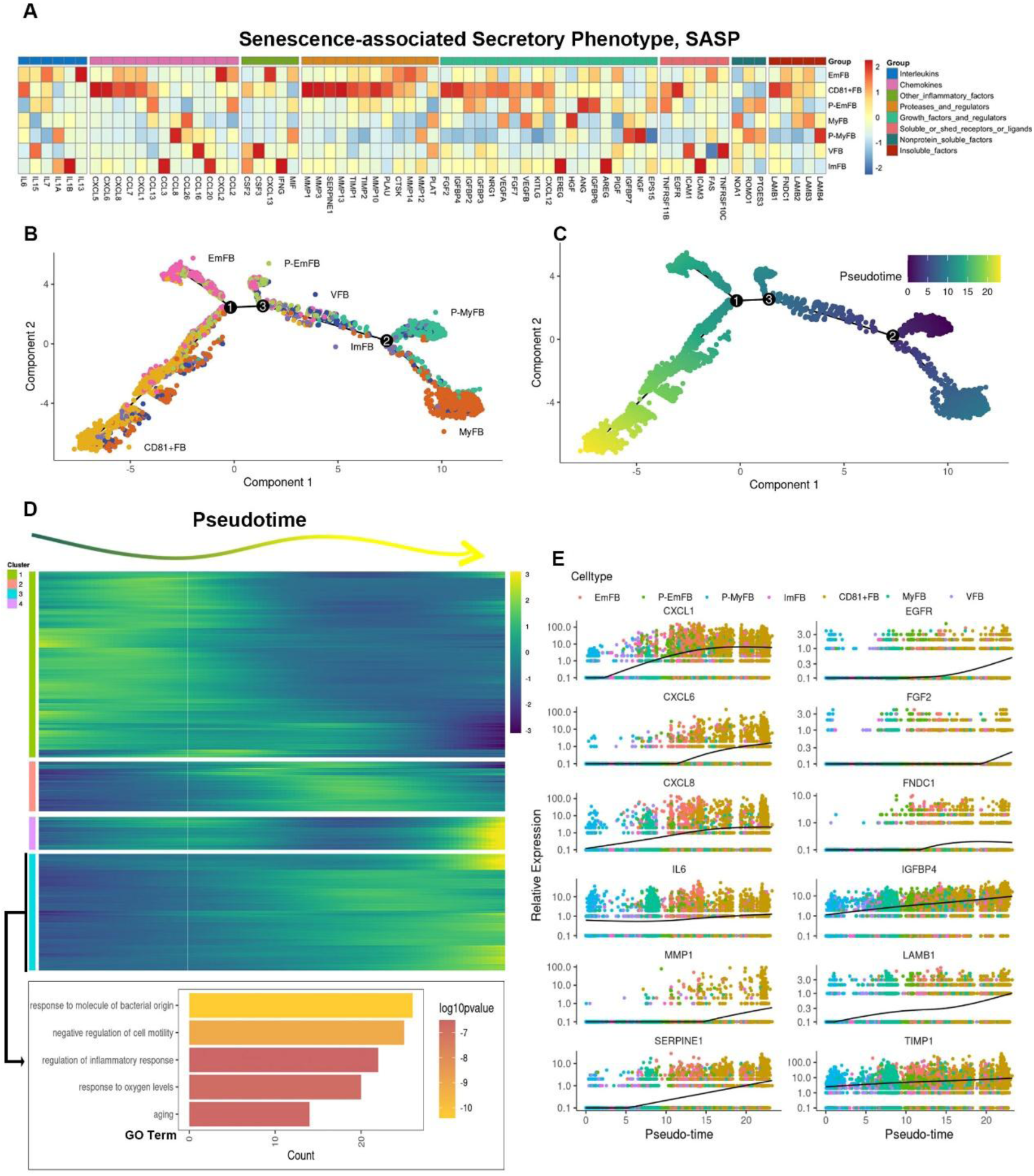
CD81^+^ gingival fibroblasts are terminally differentiational cell with high SASP expression. (**A**) Heatmap showing the relative expression for SASP genes in each fibroblast subclusters. (**B**) Trajectory reconstruction of each fibroblast subclusters. (**C**) Monocle pseudotime analysis revealing the progression of gingival fibroblast clusters. (**D**) Upper panel: Heatmap showing the scaled expression of differently expressed genes in trajectory as in (C), cataloged into four gene clusters (labels on left). Bottom panel: GO analysis of expressed genes whose expression increases as the differentiation trajectory progresses. (**E**) SASP-related genes with increased expression as the differentiation trajectory progresses.

### 2.5 CD81^+^fibroblasts indirectly sustained inflammation by recruiting neutrophils via C3/C3aR1 axis

To explore the communication between CD81^+^ fibroblasts and immune cells in periodontitis, we analyzed their interactions under diseased conditions. Our results revealed that CD81^+^ fibroblasts had the highest level of communication with immune cells, particularly neutrophils, compared to other fibroblast subgroups (Figure 5A). This suggests that CD81^+^ fibroblasts play a key role in mediating the immune response during periodontitis. Additionally, we observed a significant increase in the expression of MIF and C3 signaling pairs between CD81^+^ fibroblasts and immune cells (Figure 5B). Previous studies have demonstrated the importance of sustained neutrophil infiltration in the progression of periodontitis, with C3 known to recruit neutrophils and contribute to the formation of neutrophil extracellular traps (Ando et al., 2024; Kim et al., 2023; Song et al., 2023). Further analysis of the C3 pathway showed that the C3 receptor-ligand pair was active in the communication between CD81^+^ fibroblasts and neutrophils in both healthy and periodontitis conditions (Figure 5C), underscoring its unique role in neutrophil recruitment. Notably, CD81^+^ fibroblasts exhibited the highest expression of the C3 ligand compared to other fibroblasts subgroups, while the C3 receptor (C3aR1) was exclusively expressed by neutrophils in periodontitis (Figure 5D). We also detected higher C3 expression in human periodontitis gingiva (Figure 5E), indicating its involvement in the disease. In vitro experiments further confirmed that periodontitis gingival fibroblasts secreted higher levels of C3 protein at 30 ng/mL compared to healthy fibroblasts at about 20 ng/mL (Figure 5F), and Pg-LPS stimulation could enhance C3 secretion by gingival fibroblasts from baseline at 10 ng/mL to about 20 ng/mL (Figure 5G). Interestingly, spatial transcriptomic analysis of gingival tissue revealed that the regions expressing CD81 and SOD2, a neutrophil marker, in periodontitis overlapped in the gingival lamina propria, showing a high spatial correlation (Figure 5H). These findings suggest that CD81^+^ fibroblasts might facilitate neutrophil infiltration through the C3/C3aR1 axis, contributing to the inflammatory response in periodontitis.

**Figure 5.**
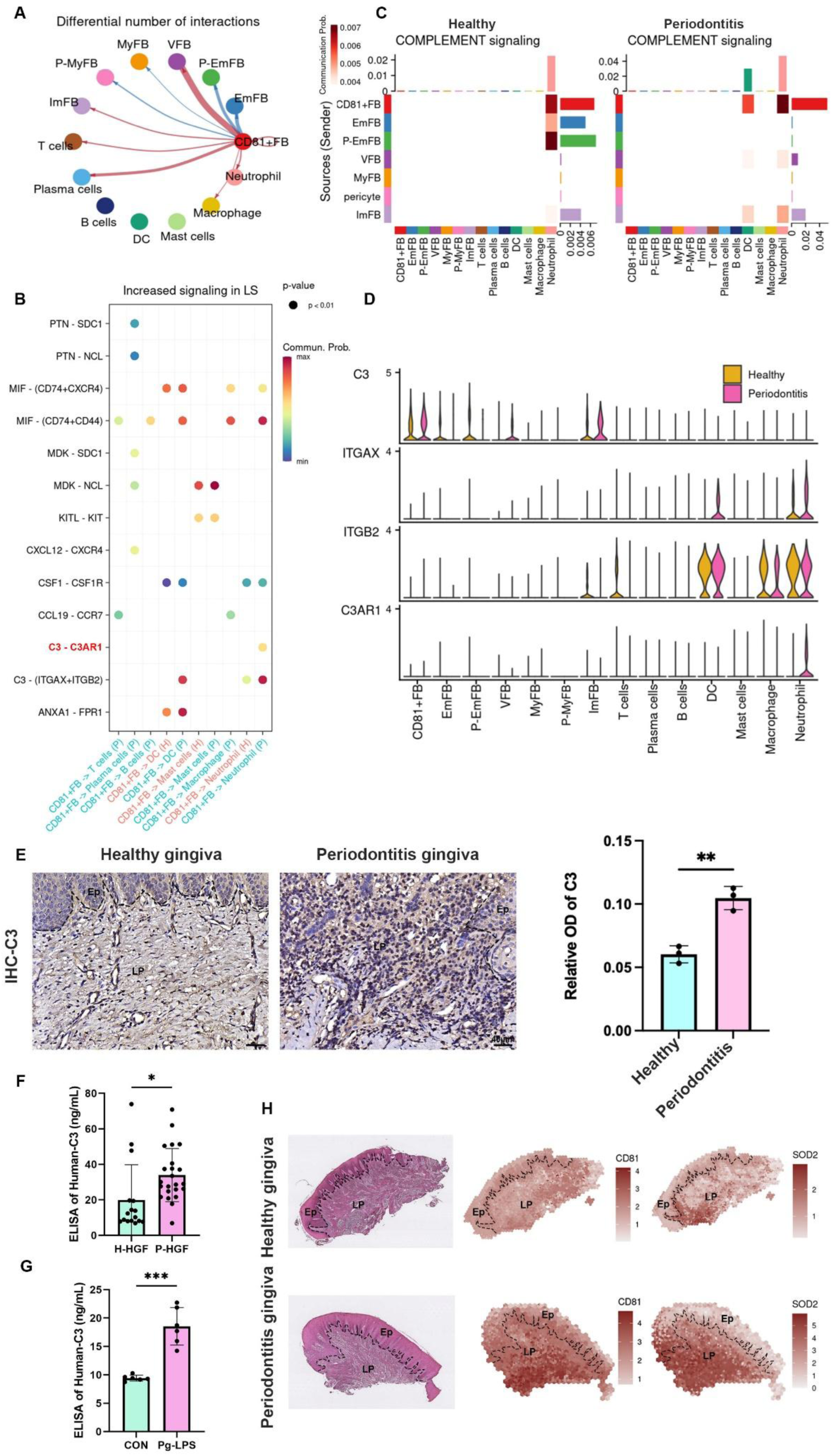
CD81^+^fibroblasts possibly recruit neutrophils via C3/C3aR1 axis. (**A**) The relative number of interactions between CD81^+^fibroblasts and other cell type in periodontitis gingiva. (**B**) Significant increased Ligand-Receptor interaction derived from CD81^+^fibroblasts. C3-C3AR1 signaling axis increased between CD81^+^ fibroblast and neutrophil especially, which was highlighted by red. (**C**) The heatmap showing the communication patterns of the Complement signaling pathway between fibroblasts and immune cell type in healthy and periodontitis gingiva. (**D**) The expression level of four representative genes in Complement signaling pathway. (**E**) Representative image of and semi-quantification of IHC staining regarding C3 in healthy and periodontitis gingiva. Scale bar=40μm, n=3. (**F**) Elisa analysis of human-C3 secretion between healthy human gingival fibroblasts (H-HGF, n= 16 samples) and periodontitis human gingival fibroblasts (P-HGF n= 23 samples). (**G**) Elisa analysis of human-C3 secretion in healthy human gingival fibroblasts with (Pg-LPS group) or without (CON group) 1 μg/mL Pg-LPS stimulated, n=6 samples. (**H**) H&E image and representative spatial mapping of CD81 and SOD2 in healthy and periodontitis gingiva from public dataset GSE152042. Co-localization in CD81 and SOD2, a neutrophil marker, was found in the periodontitis gingiva. Ep: Epithelium; LP: Lamina propria. Data are expressed as mean ± SD. *P <= 0.05, **P<= 0.01, ***P<= 0.001, ****P<= 0.0001.

### 2.6 Targeting cellular senescence in periodontitis could alleviated inflammation and bone resorption

In human periodontitis gingiva, we found that CD81^+^ fibroblasts might activate neutrophils via the C3/C3aR1 axis to exaggerate inflammation. To verify whether this mechanism exists in the LIP mouse model, we examined the expression of related markers. In the gingiva of the LIP model, P16^+^ fibroblasts, identified by p16 and Vimentin protein, comprised approximately 70% of total fibroblasts, significantly higher than the 10% observed in healthy mice (Figure 5-figure supplement 1A). Similarly, CD81^+^ fibroblasts accounted for about 30% of total fibroblasts, compared to less than 10% in the control group (Figure 5-figure supplement 1B). Immunofluorescence staining revealed co-localization of Vimentin, p16, and CD81 in LIP gingiva, indicating the presence of senescent CD81^+^ fibroblasts in the experimental periodontitis model (Figure 5-figure supplement 1C). We also observed a higher expression of C3 protein expression in the LIP group compared to controls (Figure 5-figure supplement 2A, B). Neutrophil infiltration, marked by MPO increased from approximately 10% at baseline to 40% in the inflamed gingiva of LIP mice, notably in the epithelial and lamina layers (Figure 5-figure supplement 2C, D). Further staining demonstrated that CD81^+^ C3^+^ fibroblasts constituted the majority of fibroblasts in the LIP group (Figure 5-figure supplement 2E, F). Notably, MPO^+^ neutrophils clustered around CD81^+^ cells in the lamina of the LIP model (Figure 5-figure supplement 2G). These findings in the LIP mouse model suggest that CD81^+^ fibroblasts with senescence characteristics might activate neutrophils through C3, similar to the mechanism observed in human periodontitis.

To further explore the role of senescent cells in periodontitis progression, we established a LIP mouse model treated with the senolytic drug ABT263, a Bcl2 inhibitor (Figure 6A). H&E staining revealed that the ABT263-treated group exhibited reduced inflammatory cell infiltration in the gingiva compared to the vehicle control (Figure 6B). IHC staining of senescence markers p16 and H3K9me3 showed a significant reduction in senescent cells: P16^+^ cells decreased from 20% to 8%, and H3K9me3^+^ cells from 35% to 20%, after ABT263 administration (Figure 6C, D, a and b). To assess the effect of ABT263 on eliminating CD81^+^ fibroblasts in periodontitis, immunofluorescence staining demonstrated a drop in the proportion of CD81^+^ fibroblasts from 40% to less than 20% after treatment (Figure 6E and c). Since our results suggested that CD81^+^ fibroblasts might activate neutrophil infiltration via the C3/C3aR1 axis, we next evaluated the impact of senolytic treatment on C3 secretion and neutrophil infiltration. IHC analysis revealed a slight reduction in C3 intensity in gingival tissue and a significant decrease in the number of infiltrated neutrophils after ABT263 treatment (Figure 6F, G, d and e). Finally, we observed a marked reduction in the number of osteoclasts marked by CTSK in the ABT263-treated group, decreasing from 6 cells/mm² in the vehicle group to 1 cell/mm², which suggested less bone resorbing after ABT263 treatment (Figure 6H and f). Taken together, these results suggest that senolytic treatment with ABT263 could be a potential strategy to mitigate inflammation and bone resorption in periodontitis progression.

**Figure 6.**
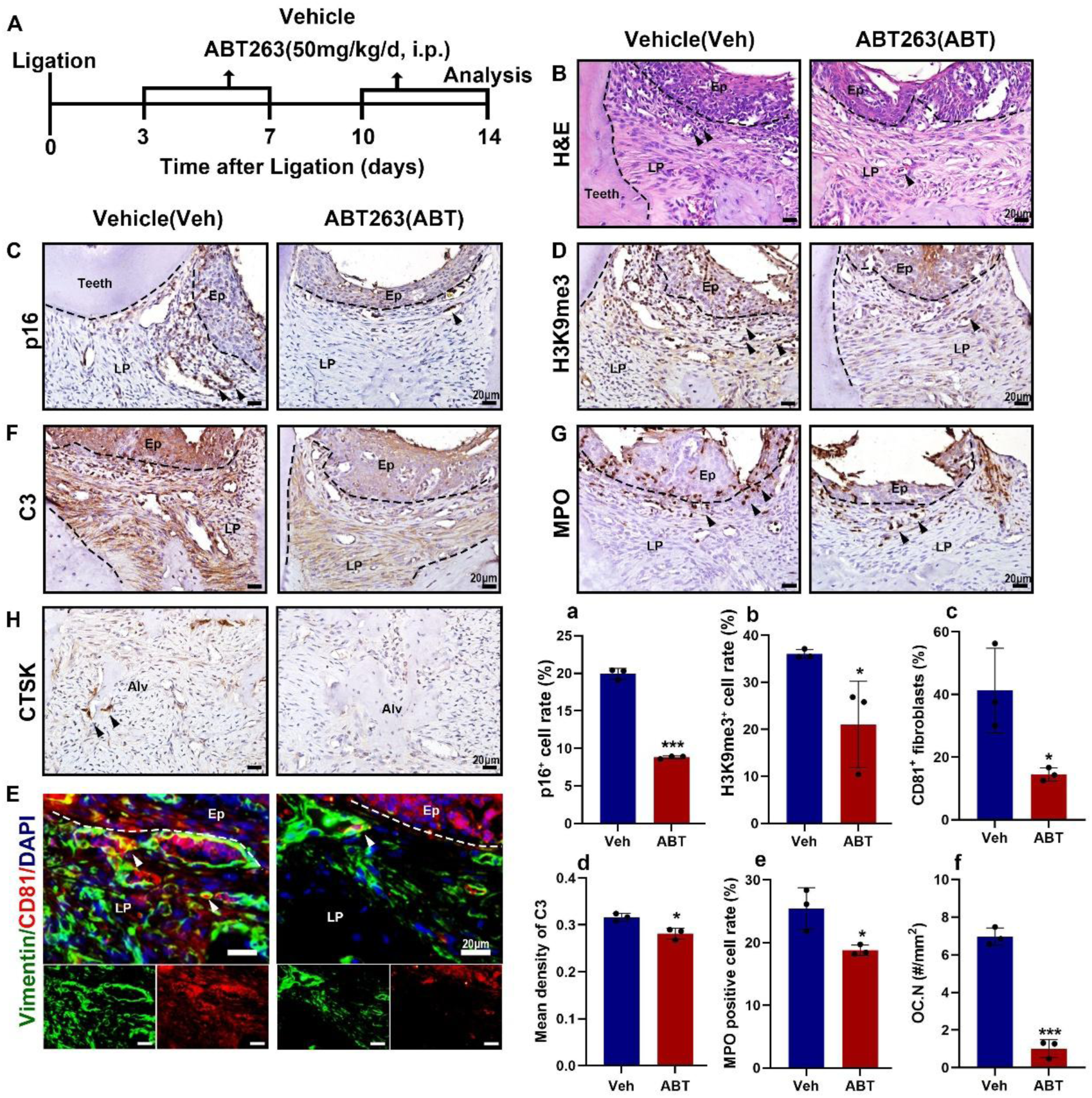
Senolytics therapy alleviates inflammation and bone resorption in LIP model. (**A**) Strategy of LIP mouse model treated by a senolytic drug Navitoclax. (**B**) Representative H&E staining image of each group, Inflammatory cells were labelled by black arrows, scale bar=20μm. (**C)** IHC staining and (**a**) semi-quantification of p16 in each group, Positive cells were labelled by black arrows, n=3, scale bar=20μm. (**D)** IHC staining and (**b**) semi-quantification of H3K29me3 in each group, Positive cells were labelled by black arrows, n=3, scale bar=20μm. (**E)** Immunofluorescence staining and **(c)** semi-quantification of CD81 (red), Vimentin (green), and nuclei (blue) in control and LIP mouse gingiva, n=3 mice, scale bar=20μm. White arrow indicates double positive cells. **(F)** IHC staining and (**d**) semi-quantification of C3 in each group, Positive cells were labelled by black arrows, n=3, scale bar=20μm. **(G)** IHC staining and (**e**) semi-quantification of MPO in each group, Positive cells were labelled by black arrows, n=3 field per group, scale bar=20μm. **(H)** IHC staining and (**f**) semi-quantification of CTSK in each group, Positive cells were labelled by black arrows, n=3 field per group, scale bar=20μm. Ep: Epithelium; LP: Lamina propria; Alv: Alveolar bone; Teeth. Data are expressed as mean ± SD. *P <= 0.05, **P<= 0.01, ***P<= 0.001, ****P<= 0.0001.

### 2.7 Metformin alleviated the inflammation and bone resorption of periodontitis via inhibiting the interaction between CD81^+^ fibroblasts and neutrophil cell

Metformin, an oral antihyperglycemic drug, has been preliminarily validated its therapeutic efficacy in periodontitis (Neves et al., 2023). However, the underlying mechanisms remain unclear. Increasing evidence suggests that metformin regulates cellular senescence, but its involvement in periodontitis-related senescence has yet to be reported (Kodali et al., 2021; Soukas, Hao, & Wu, 2019). To uncover the role of metformin in periodontitis regarding cellular senescence, we first re-analyzed sc-RNA seq data from GSE242714, which included the gingival tissue of periodontitis mice treated with metformin (Neves et al., 2023). Notably, we found that metformin treatment reduced the cellular senescence score of the periodontitis gingiva compared to untreated periodontitis gingiva (Figure 7-figure supplement 1A). Based on this, we further established a LIP mouse model and administered metformin daily for 14 days before and after the modeling time point to evaluate its effects on cellular senescence in periodontitis (Figure 7A). Micro-CT imaging revealed that delayed bone loss around the periodontal area was found following metformin administration (Figure 7B), with the higher bone volume to tissue volume ratio (BV/TV, Figure 7a) and less distance of cement-to-enamel junction to alveolar bone crest (CEJ-ABC distance, Figure 7b) compared to LIP treated with ddH_2_O group. Histological analyses further demonstrated that metformin significantly mitigated periodontitis-induced inflammatory cell infiltration (Figure 7-figure supplement 1B), and collagen degradation (Figure 7-figure supplement 1C and a), as shown by H&E and Masson staining. Additionally, metformin reversed the upregulation of p16 (Figure 7C and c), p21 (Figure 7-figure supplement 1D and b), and H3K9me3 (Figure 7-figure supplement 1E and c) in the periodontitis model. Importantly, the number of CD81^+^ fibroblasts was reduced in the LIP model after metformin administration compared to the untreated LIP group as well (Figure 7D and d). Furthermore, metformin reversed the elevated expression of C3 and MPO in periodontitis, compared to the periodontitis and ddH_2_O groups (Figure 7E, F, e, and f). In vitro, senescent fibroblasts induced by PG-LPS were treated with metformin (Figure 7-figure supplement 2A). Results showed that metformin decreased the proportion of SA-β-gal positive fibroblasts from 50% in the LPS group to 35% (Figure 7-figure supplement 2B and C). Metformin also reversed the protein expression of CD81, C3, and p16 in fibroblasts (Figure 7-figure supplement 2D). Additionally, metformin reduced the proportion of CD81/p16 and CD81/C3 double-positive gingival fibroblasts (Figure 7-figure supplement 3A-D). Collectively, these findings suggest that metformin alleviates inflammation and bone resorption in periodontitis by inhibiting the interaction between CD81^+^ fibroblasts and neutrophils, which provides a novel therapeutic strategy for periodontitis.

**Figure 7.**
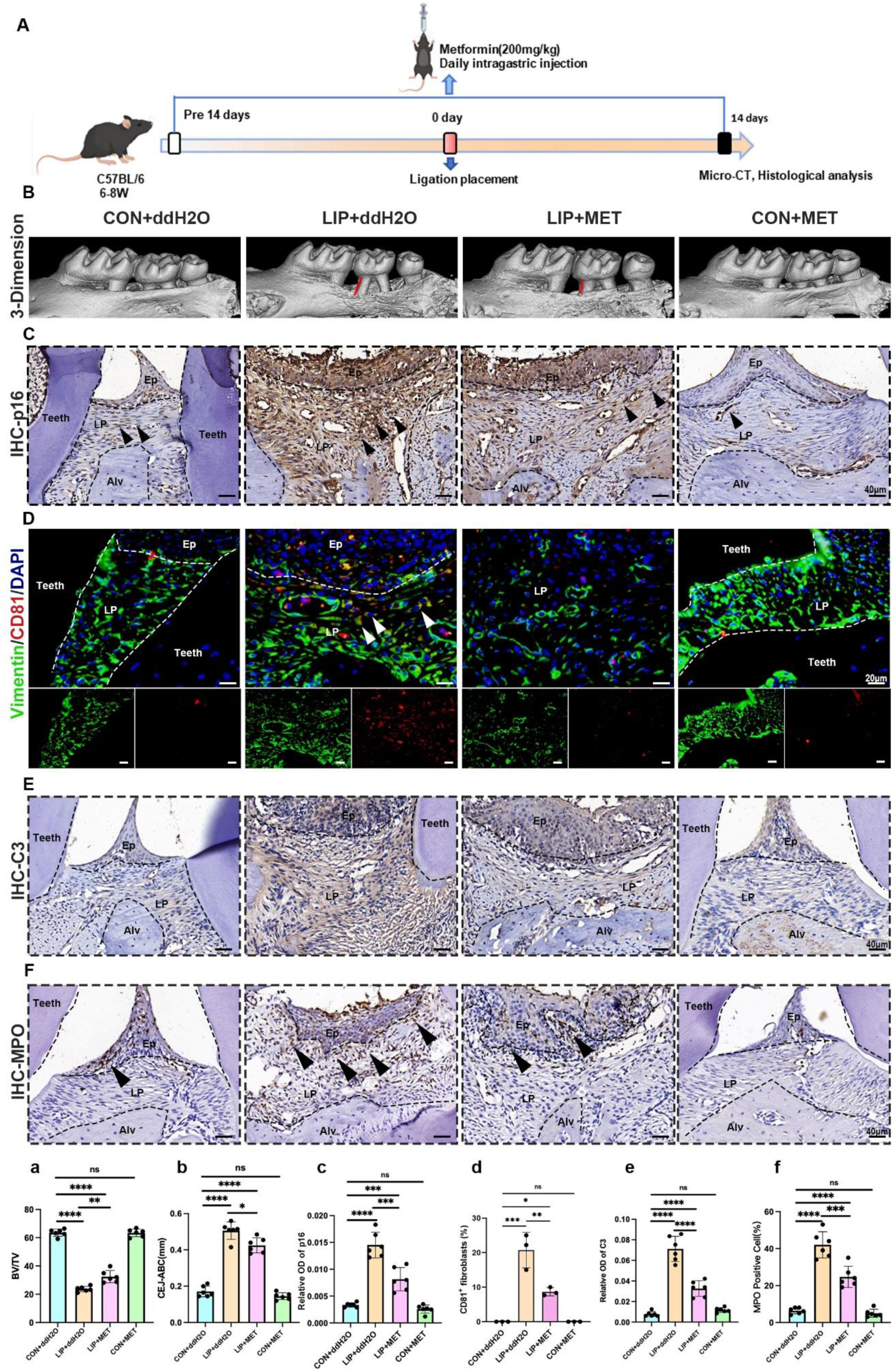
Metformin alleviates inflammation and bone resorption in LIP model via inhibiting the interaction between CD81^+^ fibroblasts and neutrophils. (**A**) Strategy of LIP mouse model treated by metformin. (**B**) 3-D visualization of the maxilla and quantified by the Bone volume/ tissue volume (BV/TV) ratio (**a**) the cement-to-enamel to alveolar bone crest, CEJ-ABC distance (**b**) indicated by red line, n= 6 mice per group. (**C)** IHC staining and semi-quantification (**c)** of p16 in each group, Positive cells were labelled by black arrows, n= 6 field per group, scale bar=40μm. (**D**) Immunofluorescence staining and **(d)** semi-quantification of CD81 positive fibroblasts in each group. Vimentin (green), CD81 (red) and nuclei (blue). White arrow indicates double positive cells, n=3, scale bar=20μm. (**E)** IHC staining and (**e)** semi-quantification of C3 in each group, n=6, scale bar=40μm. (**F)** IHC staining and (**f)** semi-quantification of MPO, a neutrophils marker, in each group, Positive cells were labelled by black arrows, n=6, scale bar=40μm. Ep: Epithelium; LP: Lamina propria; Alv: Alveolar bone; Teeth. Data are expressed as mean ± SD. *P <= 0.05, **P<= 0.01, ***P<= 0.001, ****P<= 0.0001.

## 3. Discussion

This study suggests that cellular senescence plays a role in the progression of periodontitis, and targeting cellular senescence may help alleviate the condition. We discovered that senescent gingival fibroblasts are associated with periodontitis pathology. Under continuous stimulation of LPS-PG, oxidative stress caused by ROS accelerates cell senescence in gingival fibroblasts. These senescent cells highly express CD81, which contribute to the expansion of inflammation through proinflammatory metabolic activities and factors related to SASP. Additionally, they continuously recruit neutrophils through the C3 pathway, indirectly maintaining the inflammatory response. The use of Navitoclax and Metformin can slow down the progression of periodontitis by reducing gingival cell senescence (Scheme 1).

The underlying mechanism of immune homeostasis instability and the transformation of chronic gingivitis into periodontitis has not been fully elucidated. Our findings will provide valuable insights for future studies on the pathological mechanism of periodontitis development. Gingival fibroblasts, which are essential cells in gingival connective tissue, have recently gained attention as key participants (Wielento, Lagosz-Cwik, Potempa, & Grabiec, 2023). Previous studies have reported heterogeneity in gingival fibroblasts in periodontal tissues, with four subsets significantly altered in periodontitis: Fib1.1 (CXCL1, CXCL2, CXCL13); Fib 1.2 (APCDD1, IGFBP2, MRPS6); Fib 1.3 (APOD, GSN, CFD); and Fib 1.4 (TIMP3, ASPN, COL11A1). Some of these clusters are directly associated with neutrophils and proinflammatory cytokines, suggesting that periodontal tissue immunity relies on strong matrix-neutrophil interactions within these tissues (Williams et al., 2021). Another study revealed the presence of genetic markers in a unique subgroup of gingival fibroblasts called AG fibroblasts (fibroblasts activated to guide chronic inflammation). These fibroblasts may have functional capabilities as oral immune surveillance agents and play a role in coordinating the initiation of gingival inflammation (Kondo, Gleason, Okawa, Hokugo, & Nishimura, 2023). Caetano et al. conducted a study where they mapped stromal cells from healthy and periodontitis individuals. They identified a subset of fibroblasts that expressed ARGE pro-inflammatory genes at high levels (Caetano et al., 2023). In a more recent study, the team used multiomics techniques and fluorescence in situ hybridization to demonstrate the presence of a spatially restricted population of pathogenic fibroblasts in the gingival lamina propria. These fibroblasts expressed CXCL8 and CXCL10 and were responsible for recruiting neutrophils and lymphocytes in the periodontal pocket area. Additionally, they exhibited angiogenic properties (Caetano et al., 2023). The increasing amount of data supports the role of gingival fibroblast heterogeneity in the pathological mechanism of periodontitis, particularly in immune regulation (Yin et al., 2025). However, previous studies have mainly focused on immune disorders resulting from communication between fibroblasts and immune cells, neglecting the dynamic changes of fibroblasts themselves in periodontitis pathology. In this study, we present a unique subset of fibroblasts with significantly altered gene signatures due to cell senescence, suggesting that cell senescence plays a crucial role in the heterogeneity of gingival fibroblasts. It has been recognized that low concentrations of ROS produced during chronic inflammation can indirectly cause periodontal tissue destruction (Chapple & Matthews, 2007). Recent studies have also found that repeated exposure to LPS, a component of gram-negative bacterial membranes, leads to DNA damage in various cell types, including gingival and alveolar bone cells (Aquino-Martinez et al., 2020). Cells that survive from persistent DNA damage acquire a senescent phenotype, which in turn triggers the recruitment of immune cells through dysregulation of proinflammatory cytokines. Senescent cells often overexpress IL-6, IL-1 α, IL-1 β, and IL-8, collectively referred to as SASP (Coppé et al., 2010). Our findings indicate that gingival fibroblast senescence directly promotes the development of chronic periodontitis by secreting SASP-related factors, which may explain the formation of pro-inflammatory fibroblasts and their significant impact on immune regulation. Accumulating evidence suggests that drugs can regulate the activity of SASP, as demonstrated by An et al. who showed that short-term treatment with rapamycin can reduce gingival and alveolar bone inflammation and promote the regeneration of alveolar bone in elderly mice (An et al., 2020). Additionally, Kuang et al. reported that metformin inhibits the destructive effect of H_2_O_2_ on human PDLSC, leading to a reduction in oxidative stress-induced aging (Kuang et al., 2020). Through oral administration of metformin, we have demonstrated its potential in alleviating the progression of periodontitis by delaying the senescence of gingival fibroblasts. However, further experiments are required to determine the decisive role of fibroblast senescence in the periodontitis.

CD81, a member of the tetraspanin family of proteins, could serve as a cell surface marker (Karam, Méresse, Kremer, & Daher, 2020) and a signaling pathway receptor (Oguri et al., 2020). CD81 is a major regulator of virus entry into cells and plays an important role in other pathogenic human viruses (New et al., 2021). Research on the role of CD81 has shown that it could form a complex with αV/β1 and αV/β5 integrins to activate the FAK signaling pathway (Oguri et al., 2020), which induce the interferon signaling pathway for immune response regulation (Hanagata & Li, 2011), and mediate NF-kB signaling pathway to induce IL-6 expression (Ding et al., 2019). Clinical studies have indicated a correlation between the level of CD81 in saliva and the severity of periodontitis disease (Tobon-Arroyave, Celis-Mejia, Cordoba-Hidalgo, & Isaza-Guzman, 2019), as well as its association with the regulation of aging and inflammation (Y. Jin et al., 2018). In our study, we observed that gingival fibroblasts with high CD81 expression exhibited a high enrichment of the NF-kB signaling pathway, leading to significant upregulation of IL-6 expression. The NF-kB pathway is recognized as a switch for cellular senescence, and NF-kB activation can drive cell senescence-related secretory phenotypes. Therefore, CD81 is likely to play a crucial role in regulating gingival fibroblast cell senescence. However, further investigation is needed to elucidate the specific molecular mechanism.

Finally, a link has been established between C3 from senescent fibroblasts and neutrophil infiltration in periodontitis. C3 has a strong recruitment ability for neutrophils and is crucial for the formation of neutrophil extracellular traps (NETs) (Yipp et al., 2012). Persistent neutrophil infiltration and hyperresponsiveness, including the formation of NETs, play significant roles in the development of periodontitis (Uriarte & Hajishengallis, 2023). Genetic analysis and preclinical studies have confirmed C3 as a potential pharmacological target for periodontitis treatment (Alayash et al., 2024; Hajishengallis & Chavakis, 2021). Gingival fibroblasts stimulated with IFN-γ up-regulated the expression of chemokines (CXCL9, -10, -11, CCL8), molecules involved in antigen presentation, complement component 3 (C3), and other immune response-related molecules (Ha, Jung, Choi, & Ji, 2022). Our experimental results have demonstrated that CD81^+^ gingival fibroblasts are an important source of C3. Understanding the source and mechanism of C3 complement in periodontitis is of great significance for comprehending the pathological development of the disease and can provide a new perspective for designing drug schemes.

Our study focused on identifying a specific group of gingival fibroblasts that express high levels of CD81 during the development of periodontitis. Our findings suggest that these CD81^+^ gingival fibroblasts exhibit characteristics of cellular senescence and possess strong pro-inflammatory abilities. Furthermore, we have established a connection between CD81^+^ gingival fibroblasts and the recruitment and hyperactivation of neutrophils through C3. However, further investigations are required to explore the association between CD81 and cellular senescence, as well as its potential as a therapeutic target. In conclusion, our research provides valuable insights and treatment strategies for understanding the progression of periodontitis.

## 4. Materials and methods

### 4.1 Human samples

All individuals provided written informed consent and this study was approved by the Ethics Committee of School & Hospital of Stomatology Wuhan University (WDKQ2024B01). A total of 16 participants were recruited in this study (healthy group: n=8; periodontitis group: n=8). The basic information of the included patient was listed in Supplementary Table 1. Healthy control group included patient who underwent wisdom tooth extraction or crown lengthening procedures, and inclusion criteria are as follows: 1) age 18-65 years old; 2) good general health, no systemic diseases, able to tolerate periodontal surgery; 3) no erythema, edema, bleeding and other symptoms in gingival tissue; 4) no use of nicotine-related products in recent 6 months. Periodontitis group included patients who went through pocket reduction surgeries. Inclusion criteria for patients with chronic periodontitis were as follows (Armitage, 1999): 1) age 18-65 years; 2) good general health, no systemic disease, and tolerance to periodontal surgery; 3) mild gingival tissue redness, bleeding on probing, or clinical attachment loss (CAL) ≥ 4 mm or probing depth (PD) ≥ 5 mm in non-acute inflammatory periods; 4) no use of nicotine-related products in the last 6 months. Collected gingiva were used for primary cell culture and histological analysis in this study.

### 4.2 Primary gingival fibroblast cell culture isolation and culture

Collected gingiva tissues were transported from the clinic to the laboratory in pre-cooled phosphate-buffered saline (PBS) solution and rinsed with PBS several times to remove debris. And then, the tissues were minced into small fragments at diameter of approximately 1–3 mm. The tissue pieces were digested with 2 μg/ml type II collagenase (2275GR001, BioFroxx, Germany) at 37℃ for 2 h, and collected cell precipitates were incubated for 5–7 days at 37℃ and 5% CO_2_ in DMEM high-glucose medium (DMEM, YC-2067, China) supplemented with 20% fetal bovine serum (FBS, PAN-SERATECH, South America) (J. Li et al., 2024). The primary gingival fibroblast cells that grew out of the explants were cultured and passaged. Primary gingival fibroblasts at passages four to eight were used in the following experiments. Gingival fibroblasts derived from healthy gingiva were labelled as H-HGF while those derived from periodontitis gingiva were labelled as P-HGF.

### 4.3 Pg-LPS induced human gingival fibroblasts treated with metformin

To investigate the effect of Pg-LPS on the cellular senescence of gingival fibroblasts, healthy human gingival fibroblasts (HGFs) were seeded at 5000 per well in 96 well plate and incubated in complete medium at 37 °C overnight. And then, the HGFs were stimulated by *Porphyromonas gingivalis* lipopolysaccharide (Pg-LPS InvivoGen, USA) at 0, 0.5, 1, 5 and 10 μg/mL for 24 h. At last, the samples were used for SA-β-gal staining.

To evaluate the effect of metformin on the cellular senescence of gingival fibroblasts stimulated by Pg-LPS, HGFs at 150,000 cells per ml using hemacytometer, were seeded in 3 ml plates and incubated in complete medium at 37 °C overnight. For LPS+MET group, cells were pre-treated with metformin (HY-B0627, MedChemExpress, China) at 2 mM for 24 h. And then, for LPS and LPS+MET group, cells were stimulated with Pg-LPS (InvivoGen, USA) at 1μg/mL for another 24 h according to a previous study (Sun et al., 2023). Subsequently, HGFs cells were harvested for subsequent SA-β-gal staining, western blot analysis and immunofluorescence staining.

### 4.4 Enzyme-linked immunosorbent assay (ELISA) analysis of C3

Human gingival fibroblasts were seeded in 6-well plates with 2 mL complete cell culture. When it comes to 80 or 90 % cell confluency, cells were kept in a resting state for 24 h in serum-free medium. The supernatant of cell culture was collected after centrifugal at 12,000 rpm for 20min. The concentration of C3 in cell culture supernatants was assessed by Human C3 ELISA kit (ELK1059, ELK Biotechnology, China) according to the manufacture instruction.

### 4.5 Staining for Senescence-Associated Galactosidase (SA-β-gal)

SA-β-gal staining was performed using the Senescent β-Galactosidase Staining Kit (C0602; Beyotime Biotechnology, China) according to the manufacturer’s instructions. Cell samples were incubated for 12 hours while tissue samples were incubated for 24 hours at 37°C in a CO_2_-free temperature chamber. Tissue sections were then stained by nuclear red staining solution. Positive cells were blue-stained and all cells were nuclear red-stained. Three randomized regions of interest were captured under an ordinary light microscope (DP72 microscope, Olympus, Japan) and the percentage of positive cells were counted by Image J v2.0 (NIH, Bethesda, MD, USA).

### 4.6 Ligature-induced periodontitis (LIP) mouse model treated by Senolytics or metformin C57BL/6 mice (8 weeks, male) were purchased from Hubei Provincial Center for Disease

Control and Prevention and bred in specific pathogen free animal laboratory of the School & Hospital of Stomatology, Wuhan University. The animal experiments were conducted according to the ARRIVE guidelines 2.0. Animals and approved by the Animal Research Ethics Committee at the School & Hospital of Stomatology, Wuhan University, China (No. S07922040A). The animals were housed in a SPF environment with controlled temperature/humidity with 12h light/dark cycle. To investigate the role of senescent cells in periodontitis progression, the ligature-induced periodontitis (LIP) mouse model was treated by Senolytics drug ABT263 (HY-10087, MedChemExpress, China). In brief, after anesthetics, the mice were ligated with a 5-0 silk (SA82G, ETHICON, China) between the maxillary first and second molars and knots were tied on palatal side to secure the ligature. The ligatures were examined daily to ensure that they remained in place during the experimental period. LIP mice were divided into 2 group: Vehicle and ABT263 group.

Each group included 6 mice. LIP mice were intraperitoneally injected with vehicle alone (10% DMSO + 40%PEG300 + 5% Tween-80 + 45% Saline) or with ABT263 (50mg/kg/d; HY-10087; MedChemExpress, China) as previously (S. Li et al., 2023). Three days after ligation, vehicle and ABT263 was given to mice for two cycles of 4 consecutive days, with 3 days of rest between cycles. After 14 days post ligation, mice were euthanized, and their maxilla and gingiva were collected for histological staining.

To investigate the effect of metformin on the periodontitis progression, the LIP mouse model was treated by metformin (HY-B0627, MedChemExpress, China). Mice were allocated into four group: CON+ddH_2_O group, LIP+ ddH_2_O group, LIP+MET group and CON+MET group, each group included 6 mice. LIP+MET group and CON+MET group were treated with 200 mg/kg metformin while CON+ddH_2_O group and LIP+ddH_2_O were treated with the distilled water as the control. Metformin or ddH_2_O were given by intragastric administration once a day for 14 days before LIP model establishment. On the fifteenth day after intragastric administration, LIP+ ddH_2_O group and LIP+MET group were ligated with a 5-0 silk between the maxillary left first and second molars and knots were tied on palatal side to secure the ligature. A second set of controls included mice that were not treated with ligatures on either side. Metformin or ddH_2_O were given once a day for another 14 days. At the end of the time frame, mice were euthanized and their maxilla and gums were collected for micro-CT and histological analysis.

### 4.7 Micro-tomographic (Micro-CT) scanning and analysis

Micro-CT scanning was performed using Bruker Micro-CT SkyScan1276 (Konitich, Germany). The region of interest (ROI) was established in a three-dimensional (3-D) scope: vertically, starting from 0.2 mm apical to the cemental enamel junction (CEJ) of the second molar (2nd M), extending towards the root apical to get a span of 0.5 mm; mesiodistally, ranging from the most mesial aspect of the CEJ of the first molar (1st M) to the root furcation of the third molar (3rd M); buccolingually and lingually, ranging around the root furcation of the 2nd M within a span of 1.5 mm. The ratio of bone volume to total volume (BV/TV) was calculated based on this ROI. The distances between the CEJ to the alveolar bone crest (CEJ-ABC) were measured at the 2nd M. The 3-D reconstruction, calculation and measurements were conducted using the CTAn software (version 1.18.8.0, SkyScan, Germany). All measurements were repeated for three times with 6 mice per group, with the average value of the bilateral maxillary alveolar bone taken as one sample for statistical analysis.

### 4.8 Protein extraction and western blot

Protein extracted from mice samples or primary gingival fibroblasts were dissolved in 80 μL of RIPA buffer to extract total protein, supplemented with protease and 1% phosphatase inhibitors. All samples were quantified and normalized using a protein assay kit known as bicinchoninic acid (Thermo Fisher Scientific, Waltham, MA, United States). Following a 10-minute heat treatment at 95 ℃, the samples underwent sodium dodecyl sulfate-polyacrylamide gel electrophoresis for separation and were then transferred to a polyvinylidene fluoride membrane (Millipore). The membrane was blocked using the primary antibody-blocking solution and then incubated overnight at 4 ℃ with primary antibodies against p16 (10883-1-AP, Proteintech, China), CD81 (66866-1-IG, Proteintech, China), β-ACTIN (66009-1-Ig, Proteintech, China), C3 (21337-1-AP, Proteintech, China),and GAPDH (PMK052S, Biopm, China). Subsequently, the membrane was treated with horseradish peroxidase (HRP) conjugated secondary antibodies at 37 ℃ for 1 h. Visualization of signals were conducted using a Ultrasensitive ECL Detection Kit (Thermo Fisher Scientific, Waltham, MA, United States) with the ChemiDoc MP Imaging Systems (Bio-Rad, USA). Protein levels were normalized to β-ACTIN or GAPDH using Image J analysis software.

### 4.9 RNA extraction and RT-qPCR

To extract total RNA, the Trizol reagent and standard collection procedure were utilized. Total RNA concentration was measured using a Nanodrop2000 instrument (Thermo Fisher Scientific, Waltham, MA, United States). According to the guidelines provided by the manufacturer, the total RNA was subjected to reverse transcription into cDNA using the HiScript II Q RT SuperMix (Vazyme). The amplification reaction was performed using ChamQ SYBR qPCR Master Mix (Vazyme) in the QuantStudio 6 Flex System (Thermo Fisher Scientific, Waltham, MA, United States). The primers for the experiment were bought from Sangon Biotech Co., Ltd. The results were analyzed using the 2^−ΔΔCt^ method, with normalization to β-actin and calibration to the control group. The forward and reverse primer sequences of the target genes used in the experiment can be found in Supplementary Table 2.

### 4.10 Histological analysis

The human gingiva samples were kept in 4% paraformaldehyde for 24 h, followed by dehydrated, fixed in paraffin or optimal cutting temperature compound. The mice maxilla with gingival tissues were kept in 4% paraformaldehyde for 24 h, followed by 4 weeks of decalcification with 15% EDTA at pH 7.4. The decalcifying solution underwent replacement every 2 days. Tissues were then dehydrated, fixed in paraffin and sectioned. The sections were stained by hematoxylin and eosin (H&E), Masson, immunohistochemical and immunofluorescence staining. Immunohistochemical and immunofluorescence staining were performed According to the manufacturer’s instructions (MXB biotechnologies, Fuzhou, China). The primary antibodies used for immunohistochemistry included p16 (1:1000; Cat: 10883-1-Ap, Proteintech, China), p21 (1:200, Cat: 10355-1-AP, Proteintech, China), H3K9me3 (1:1000. Cat: M1112-3, HUABio, China), C3 (1:200, Cat: 21337-1-AP, Proteintech, China), MPO (1:200, Cat: Ab208670, Abcam) and CTSK (1:200, Cat: 121071, Proteintech, China). Immunofluorescence (IF) staining was performed with the antibodies of CD81 (1:1000, Cat: 10883-1-AP, Proteintech, China), Vimentin (1:200, Cat: A19607, ABclonal, China), p16, C3 and MPO as previously described. In immunohistochemical staining (IHC), 3,3-diaminobenzidine tetrahydrochloride (Zhongshan Biotechnology, Ltd, China) was utilized for visualization. For double IF staining, anti-mouse, and rabbit secondary antibodies had Cy3 red and 488 nm green fluorescent markers (ABclonal, China). For triple IF staining, the nucleus of cells in tissues was stained using DAPI (Zhongshan Biotechnology, Ltd, China). The stained sections were examined and captured using an Olympus DP72 microscope (Olympus Corporation, Japanese). For semi-quantification of protein expressions, the mean optical density (MOD) of positive stains was measured using the imageJ2 software (version: 2.14.0, National Institutes of Health, Bethesda, MD). For the semi-quantification of Masson’s trichrome, the collagen volume fractions (stained blue) for individual sections were measured using ImageJ2 software.

### 4.11 Bulk RNA sequencing

For bulk RNA sequencing, RNA was extracted using the methods outlined in the qRT-PCR protocol. The total RNA was then sent to the Analysis and Testing Center at the Institute of Hydrobiology, Chinese Academy of Sciences (Wuhan, China) for quality control, library preparation, and sequencing on the Illumina platform. We utilized the Illumina TruSeq RNA library preparation kit, which generated libraries with insert fragment lengths of approximately 400-500 bp. The resulting fastq reads were aligned to the mouse genome (GRCm38) using a dedicated RNA-seq aligner. We filtered the raw data quality using Trimmomatic (version 0.36) (Bolger, Lohse, & Usadel, 2014). The filtered reads were subsequently aligned to the reference genome with HISAT2 (version 2.2.1), and the aligned reads were quantified using StringTie (Pertea, Kim, Pertea, Leek, & Salzberg, 2016). The average mapping rate for each sample exceeded 90%, with sequencing depths ranging from 30M to 40M reads.

We employed DESeq2 (version 1.34.0) to identify differentially expressed gene sets, applying thresholds of |log2 (fold change)| > 1 and a significance level of P < 0.05 (Love, Huber, & Anders, 2014). The selected differentially expressed genes were then subjected to Gene Ontology (GO) enrichment analysis. Additionally, Gene Set Enrichment Analysis (GSEA) was performed using GSEA_Linux_4.1.0 to identify relevant pathways (Subramanian et al., 2005). Significant gene sets were determined based on three criteria: P < 0.05, false discovery rate (FDR) < 0.25, and an absolute normalized enrichment score (NES) > 1.

### 4.12 Single-cell RNA sequencing analysis

Single-cell RNA transcriptome including GSE164241, GSE152042 and GSE242714 were obtained from the GEO dataset. GSE164241 contained 70407 cells from 13 healthy samples and 8 periodontitis samples (Williams et al., 2021). GSE152042 contained 12379 cells from 2 healthy samples ,1 periodontitis sample (mild) and 1 periodontitis sample (severe) (Caetano et al., 2021). GSE242714 contained 6473 cells from the control mice and LIP mice samples, which were put on either water or Metformin samples (n=5 group) (Neves et al., 2023). As for scRNA-seq, the “Seurat4.4.0” package was applied to integrate different samples with CCA (cross-dataset normalization) method, GSE164241 cell profiles were filtered scriteria of Feature_RNA> 200 & nFeature_RNA <5000 & MT_percent <10 & nCount_RNA<25000 & nCount_RNA > 1000, then GSE152042 cell profiles were filtered scriteria of Feature_RNA> 500 & nFeature_RNA <6000 & MT_percent <20, and GSE242714 cell profiles were filtered Feature_RNA> 300 & nFeature_RNA <5000 & MT_percent <15 & nCount_RNA<25000 & nCount_RNA > 500 , then those data were further normalized using the “LogNormalize”method, and the unique gene markers in each group were identified with the “FindMarkers“function.“UMAP” was used to display the cell distribution. The function “Addmodulescore” was used to reflect differences in biological processes in different cell populations.

### 4.13 Fibroblast cell re-clustering analysis

Fibroblast clusters from GSE164241 were reanalysed and were then re-normalised by calling the ‘NormalizeData’ function to account for the reduction in cell numbers subsequent to subsetting the data. The top 2000 most variable features across the dataset were then identified using the ‘FindVariableFeatures’. These variable features were subsequently used to inform clustering by passing them into the ‘RunPCA’ command. Via ‘Elbowplot’, we identified that the first eight principle components should be used for downstream clustering when invoking the ‘FindNeighbors’ and ‘RunUMAP’, as detailed above.

### 4.14 Gene function enrichment analysis

GO analysis was performed using ‘Enrichr’ on the top 200 differentially expressed genes (adjusted p-value < 0.05 by Wilcoxon Rank Sum test) (Kuleshov et al., 2016). GO terms shown are enriched at FDR < 0.05. The enrichment analysis between different fibroblast subsets in scRNAseq was performed by ‘Metascape’ and further drawn by the ‘ggplot2’ package. Four methods including ‘ssGSEA’, ‘AUCell,’ ‘UCell,’and ‘singscore’ were used for enrichment analysis between different clusters. Images were further drawn by “irGSEA”. GSEA was applied to validate the result on RNA-seq with default settings (1000 permutations for gene sets, Signal2Noise metric for ranking genes).

### 4.15 “Cellular senescence” and “Senescence-associated secretory phenotypes” gene set

The cellular senescence gene set, which was used to reflect the degree of cellular senescence, has been validated across species in a variety of cell lines and multiple sequencing data including scRNA-seq, bulk RNA-seq, etc. It has better validation efficiency than previously known gene sets associated with cell senescence (Saul et al., 2022). SASP includes several soluble and insoluble factor families. These factors can affect surrounding cells by activating various cell surface receptors and corresponding signal transduction pathways that may lead to a variety of pathologies. SASP factors can be divided globally into the following main categories: soluble signal transduction factors (interleukins, chemokines and growth factors), secreted proteases and secreted insoluble protein/extracellular matrix (ECM) components (Coppé et al., 2010).

### 4.16 Metabolism pathway analysis

The ‘scMetabolism’ package was used to quantify the metabolism activity at the scRNA-seq dataset. Seventy-eight metabolism pathways in KEGG were included in the package. The pathways were further used to evaluate the metabolism activity at the single-cell resolution (Wu et al., 2022).

### 4.17 Pseudotime trajectory analysis

We applied the single-cell trajectories analysis utilizing Monocle2 using the DDR-Tree and default parameter. Before Monocle analysis, we selected marker genes from the Seurat clustering result and raw expression counts of the cell passed filtering. Based on the pseudotime analysis, branch expression analysis modeling (BEAM Analysis) was applied for branch fate determined gene analysis (Qiu et al., 2017).

### 4.18 Cell-cell communication analysis

The cell-cell communication was measured by quantification of ligand-receptor pairs among different cell types. Gene expression matrices and metadata with major cell annotations were used as input for the CellChat package (v1.6.1) (S. Jin et al., 2021).

### 4.19 Spatial transcriptomics data analysis

Spatial Transcriptomics slides were printed with two identical capture areas from 1 healthy sample and 1 periodontitis sample (Caetano et al., 2023). The capture of gene expression information for ST slides was performed by the Visium Spatial platform of 10x Genomics through the use of spatially barcoded mRNA-binding oligonucleotides in the default protocol. Raw UMI counts spot matrices, imaging data, spot-image coordinates, and scale factors were imported into R using the Seurat package (versions 4.2.2). Normalization across spots was performed with the ‘LogVMR’ function. Dimensionality reduction and clustering were performed with independent component analysis (PCA) at resolution 1 with the first 30 PCs. Signature scoring derived from scRNA-seq or ST signatures was performed with the ‘AddModuleScore’ function with default parameters in Seurat. Spatial feature expression plots were generated with the SpatialFeaturePlot function in Seurat (versions 3.2.1). To further increase data resolution at a subspot level, we applied the BayesSpace package (Zhao et al., 2021).

### 4.20 Statistical analysis

GraphPad Prism software (version 6.0, USA) was used for statistical analyses. Data were presented as the mean and standard deviation (SD) in all graphs. Data were analyzed using the unpaired Student’s t-test in order to compare group pairs or ANOVA for multiple group comparisons. Statistical significance was set at p < 0.05.

## Acknowledgements

Thanks to all clinical participants for their contribution. We Would like to thank Dr. Zhixian Qiao and Xiaocui Chai at The Analysis and Testing Center of Institute of Hydrobiology, Chinese Academy of Sciences for their assistance with RNA-seq and data analysis.

## Funding

The National Natural Science Foundation of China (No.32370816) and Research Project of School and Hospital of Stomatology Wuhan University (No.ZW202403) for Haibin Xia. Undergraduate Training Programs for Innovation and Entrepreneurship of Wuhan University, (No.S202510486507) for Min Wang. The funders had no role in study design, data collection, and interpretation, or the decision to submit the work for publication.

### Author contributions

**Liangliang Fu**: Conceptualization, Investigation, Formal analysis, Supervision, Project administration, Methodology, Writing–original draft; **Chenghu Yin**: Conceptualization, Investigation, Formal analysis, Software, Methodology, Validation, Writing–original draft; **Qin Zhao:** Methodology, Resources, Supervision; **Shuling Guo:** Software, Visualization; **Wenjun Shao:** Data curation, Software; **Ting Xia:** Methodology, Software; **Quan Sun:** Methodology, Visualization; **Liangwen Chen:** Resources, Visualization; **Jinghan Li**: Methodology, Software, Validation. **Min Wang:** Funding acquisition, Project administration, Writing–review and editing; **Haibin Xia**: Conceptualization, Funding acquisition, Project administration, Writing–review and editing.

### Ethics Statement

Human subjects: All individuals provided written informed consent and this study was supported by the Ethics Committee of School & Hospital China Hospital of Stomatology Wuhan University (WDKQ2024B01).

The animal tests in this study adhere to the guidelines established by the Animal Research Ethics Committee at the School & Hospital of Stomatology, Wuhan University, China. The Ethics Committee approved the Animal Use research with protocol number No. S07922040A.

### Data availability

Upon a reasonable request, the corresponding author is able to furnish all the essential data that backs up the discoveries of this research. Single-cell RNA-sequencing data obtained in this study are provided in NIH Gene Expression Omnibus (GSE164241, GSE152042 and GSE242714):

---

**Figure 1-figure supplement 1.**
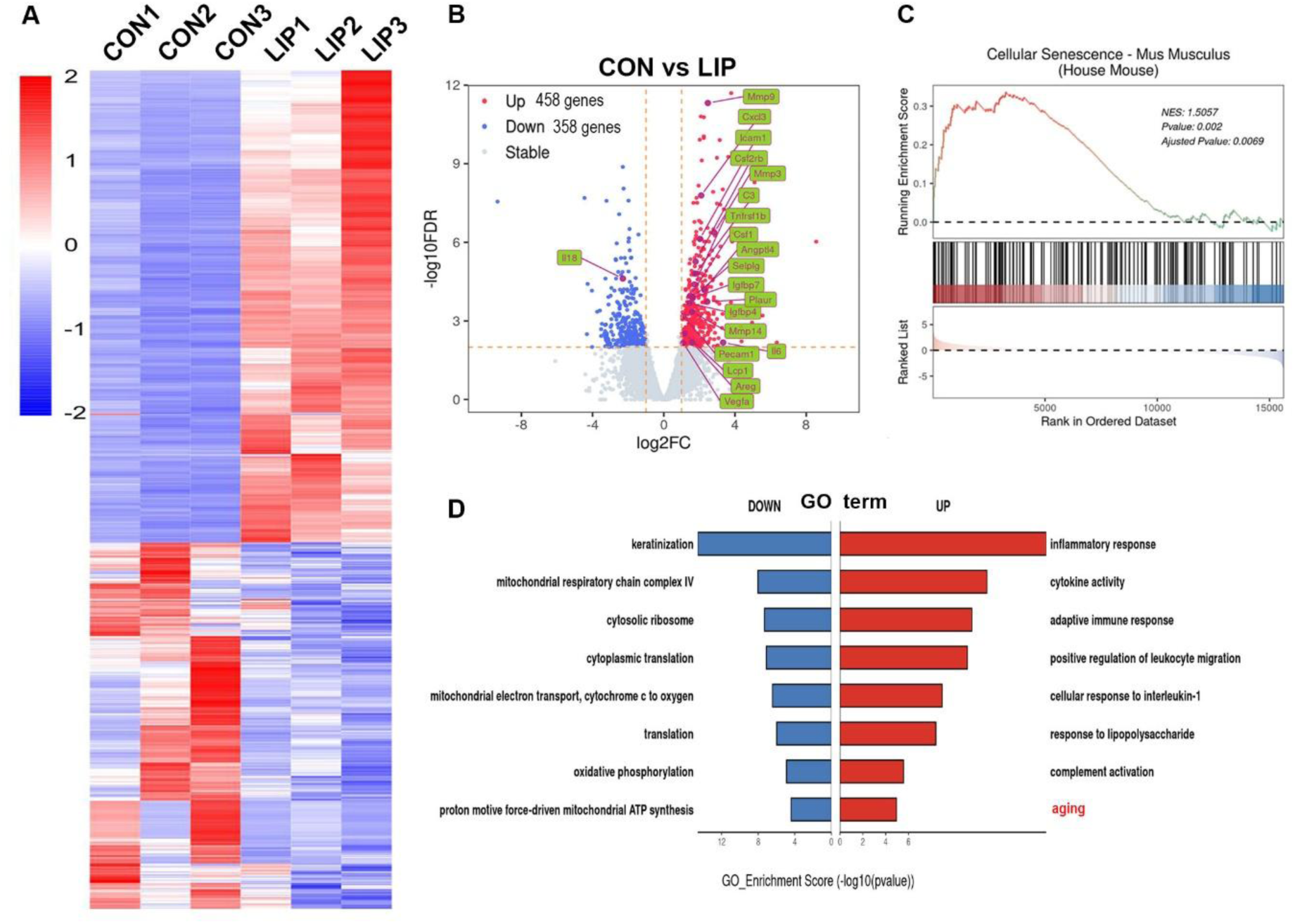
Bulk RNA-seq analysis of ligature-induced periodontitis mice model. (**A**) Heatmap and **(B)** Volcano plots of differentially expressed genes in mouse gingiva at LIP 7D compared to the CON (n=3 samples each group). Representative senescence-related genes are indicated as green. blue dots indicate differentially down-regulated genes; red dots indicate differentially up-regulated genes. Significantly different expression genes with | log2FC | > 1 and FDR < 0.05. (**C**) GSEA enrichment analysis of cellular senescence gene sets in mouse gingiva at LIP 7D compared to the CON. (**D**) GO enrichment analysis with upregulated (red) and downregulated (blue) genes shown in **(A)**. Aging biological process was significantly enriched and highlighted by red.

**Figure 1-figure supplement 2.**
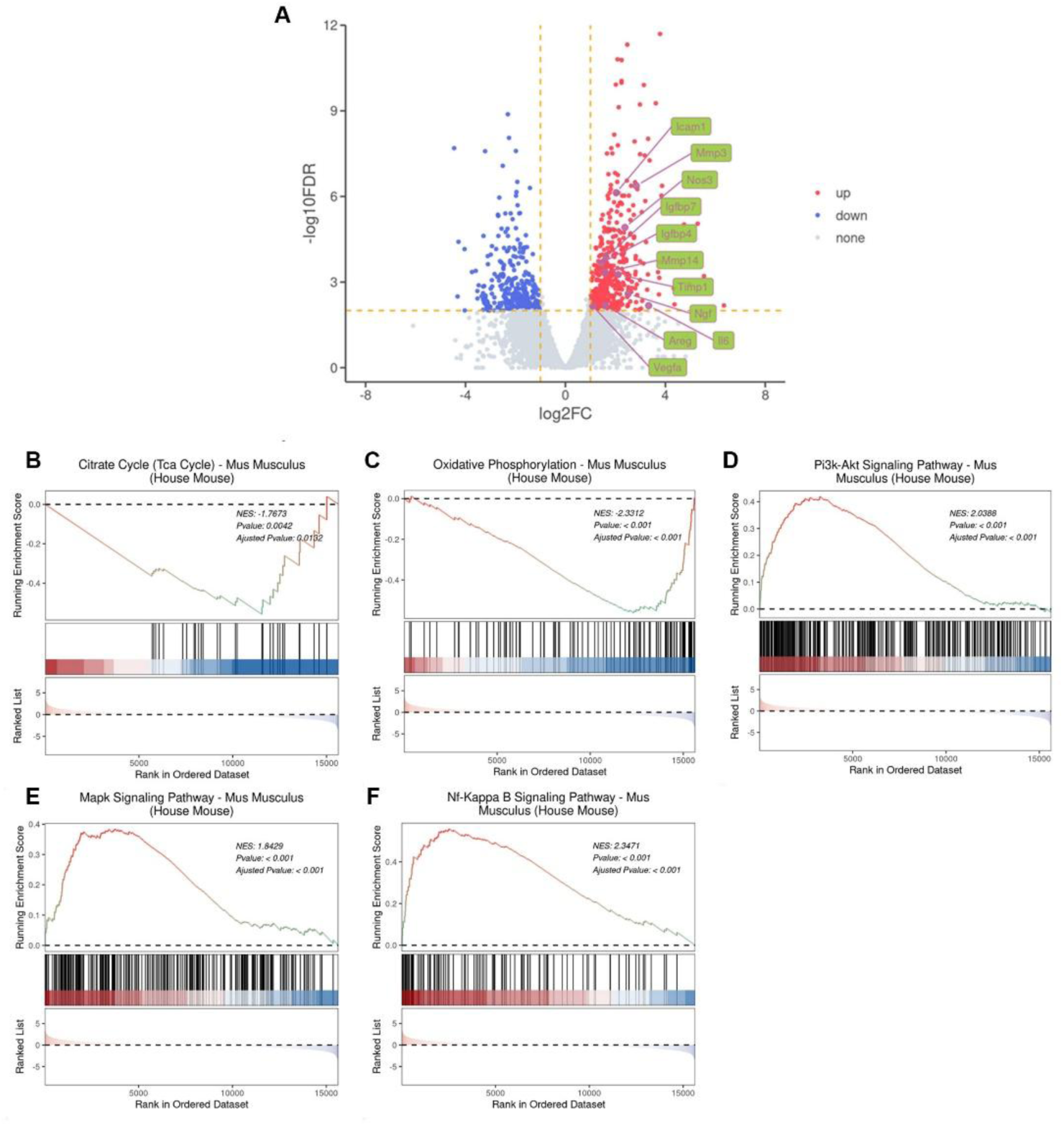
Bulk RNA-seq analysis of ligature-induced periodontitis mice model in regard to cellular senescence. **(A)** Volcano plots of differentially expressed genes in mouse gingiva at LIP 7D compared to the control (CON). Representative senescence-associated secretory phenotypes (SASP) genes are indicated as green. **(B and C)** GSEA enrichment analysis of Citrate Cycle and Oxidative Phosphorylation gene sets in mouse gingiva at LIP 7D compared to the CON, which indicated mitochondrial dysfunction in periodontitis. **(D-F)** GSEA enrichment analysis of Pi3k-Akt, Mapk and Nf-Kappa B signaling pathway gene sets in mouse gingiva at LIP 7D compared to the CON, which indicated senescence-associated signaling pathway were activated in periodontitis.

**Figure 2-figure supplement 1.**
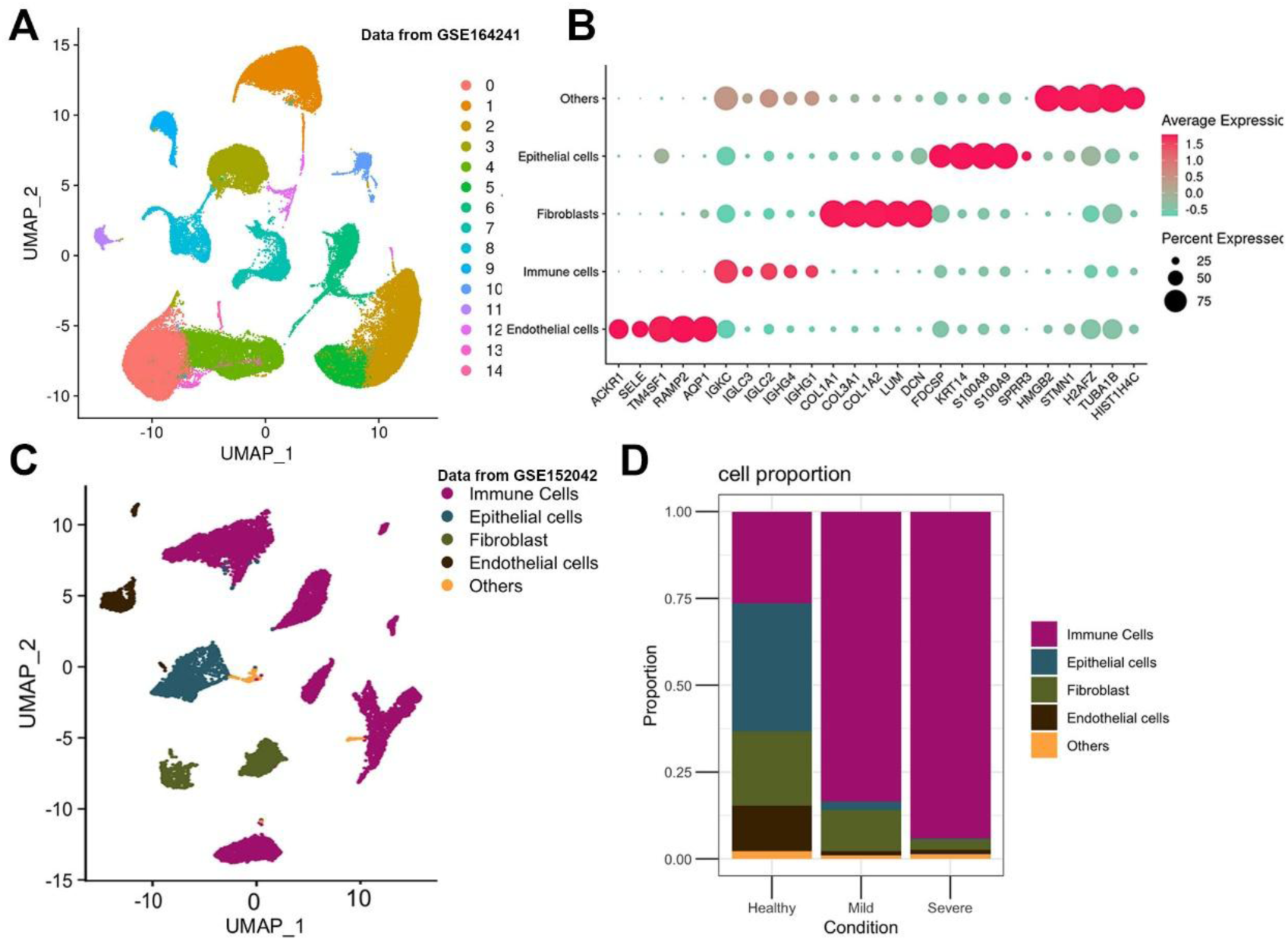
**(A)** UMAP diagram illustrated the cell clusters of GSE164241. **(B)** Marker genes of each cell were shown in the dot plot. **(C)** UMAP diagram and single-cell annotation of cells clusters from GSE152042. **(D)** Histogram of gingival tissue cell ratio in healthy, mild and severe periodontitis patients from GSE152042.

**Figure 2-figure supplement 2.**
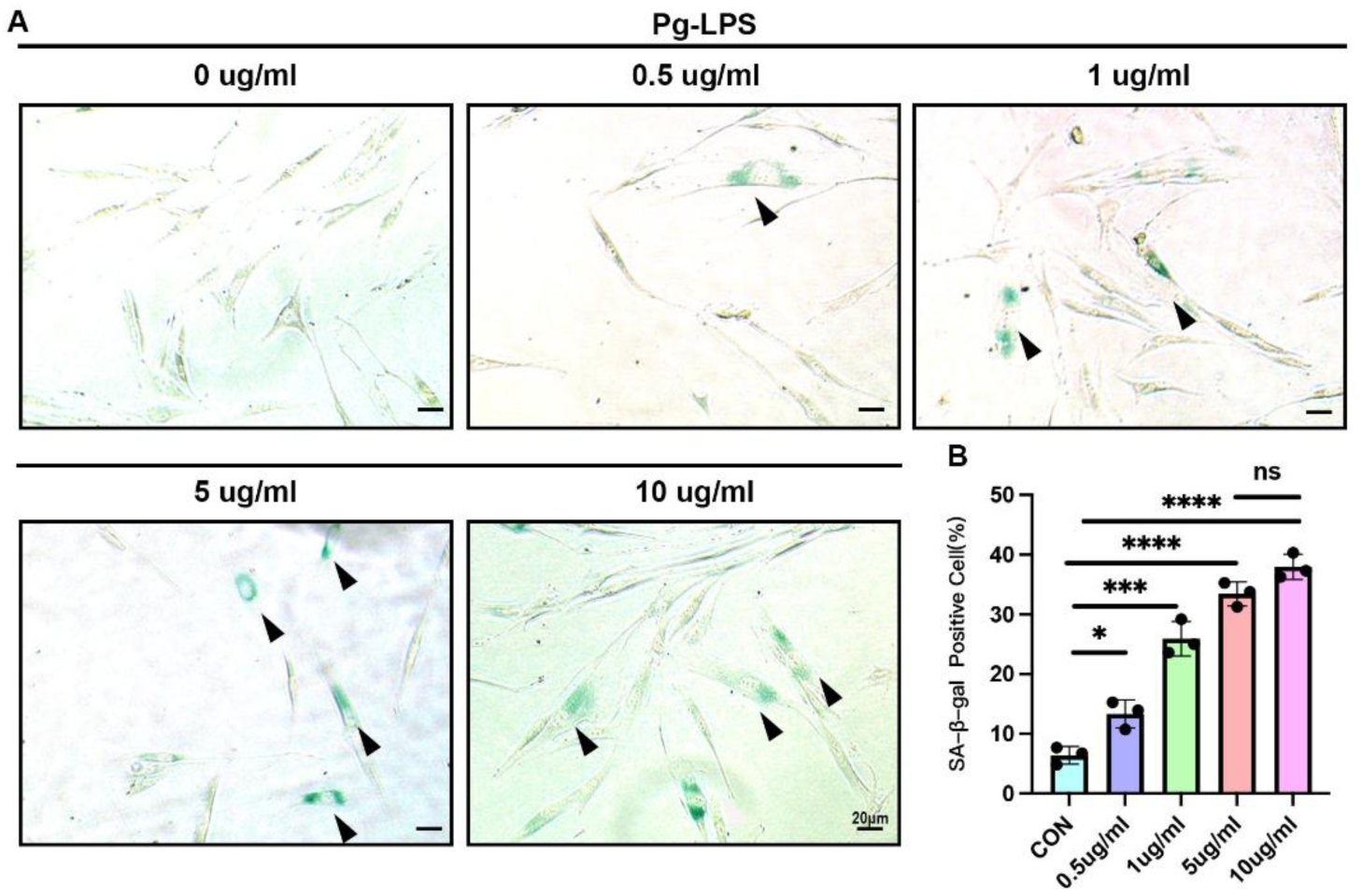
(**A**) SA-β-gal staining and **(B)** semi-quantification of human gingival fibroblasts stimulated by different concentrations of Pg-LPS (n=3). Black arrow indicates SA-β-gal positive cells. Data are expressed as mean ± SD. *P <= 0.05, **P<= 0.01, ***P<= 0.001, ****P<= 0.0001.

**Figure 4-figure supplement 1.**
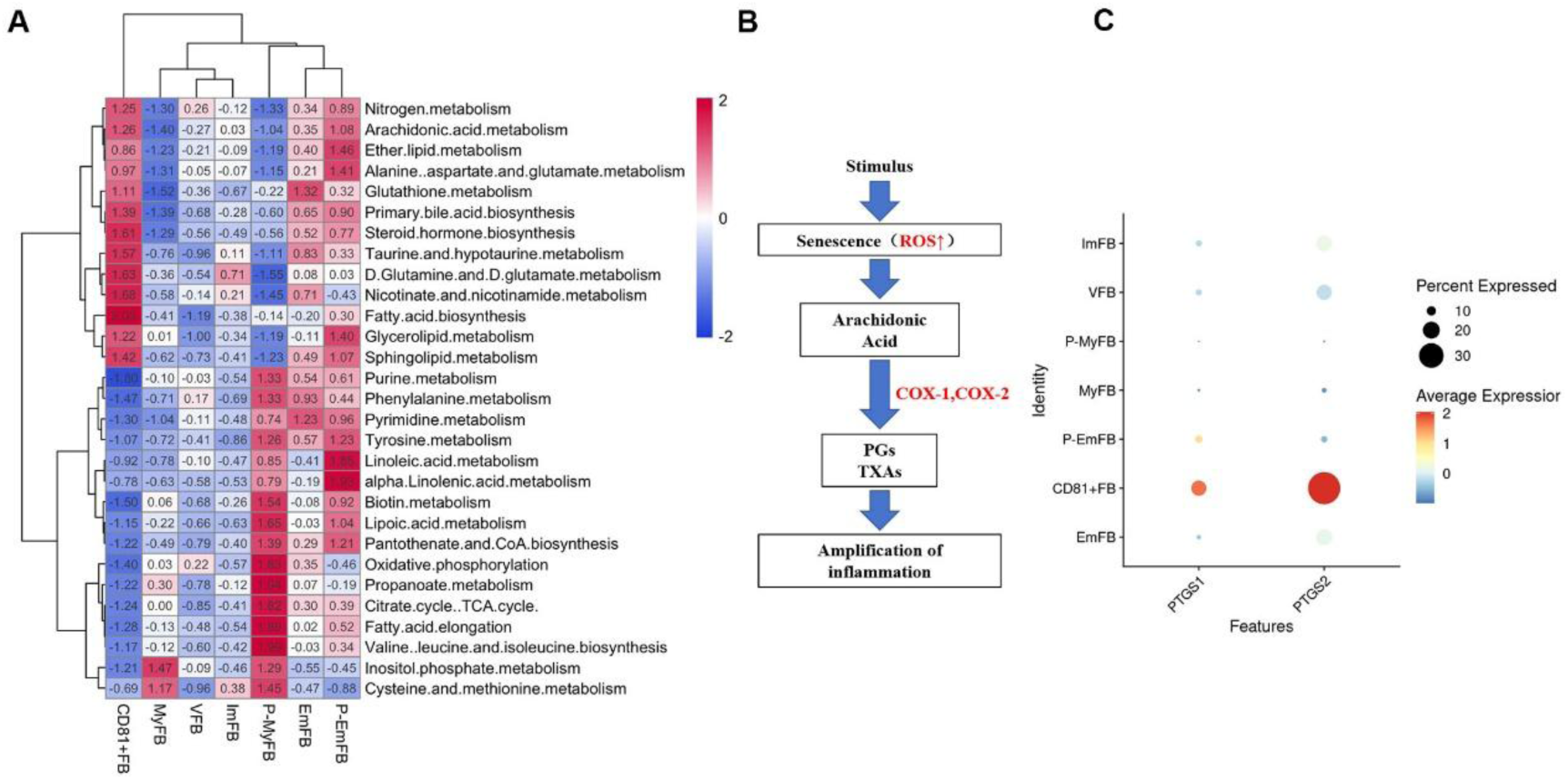
(**A**) The heatmap representing metabolic pathways in each fibroblast subclusters, which indicated fatty acid biosynthesis, arachidonic acid metabolism, and steroid biosynthesis were significantly upregulated in CD81^+^ fibroblasts. (**B**) The flow chart representing the metabolism of arachidonic acid, which could be converted into prostaglandins (PGs) and Thromboxane As (TXAs) by COX-1 or COX-2. (**C**) The dot plot representing that PTGS1 gene (encoding COX1 protein) and PTGS2 gene (encoding COX2 protein) are significantly higher in CD81^+^ gingival fibroblasts compared to other fibroblasts subclusters.

**Figure 5-figure supplement 1.**
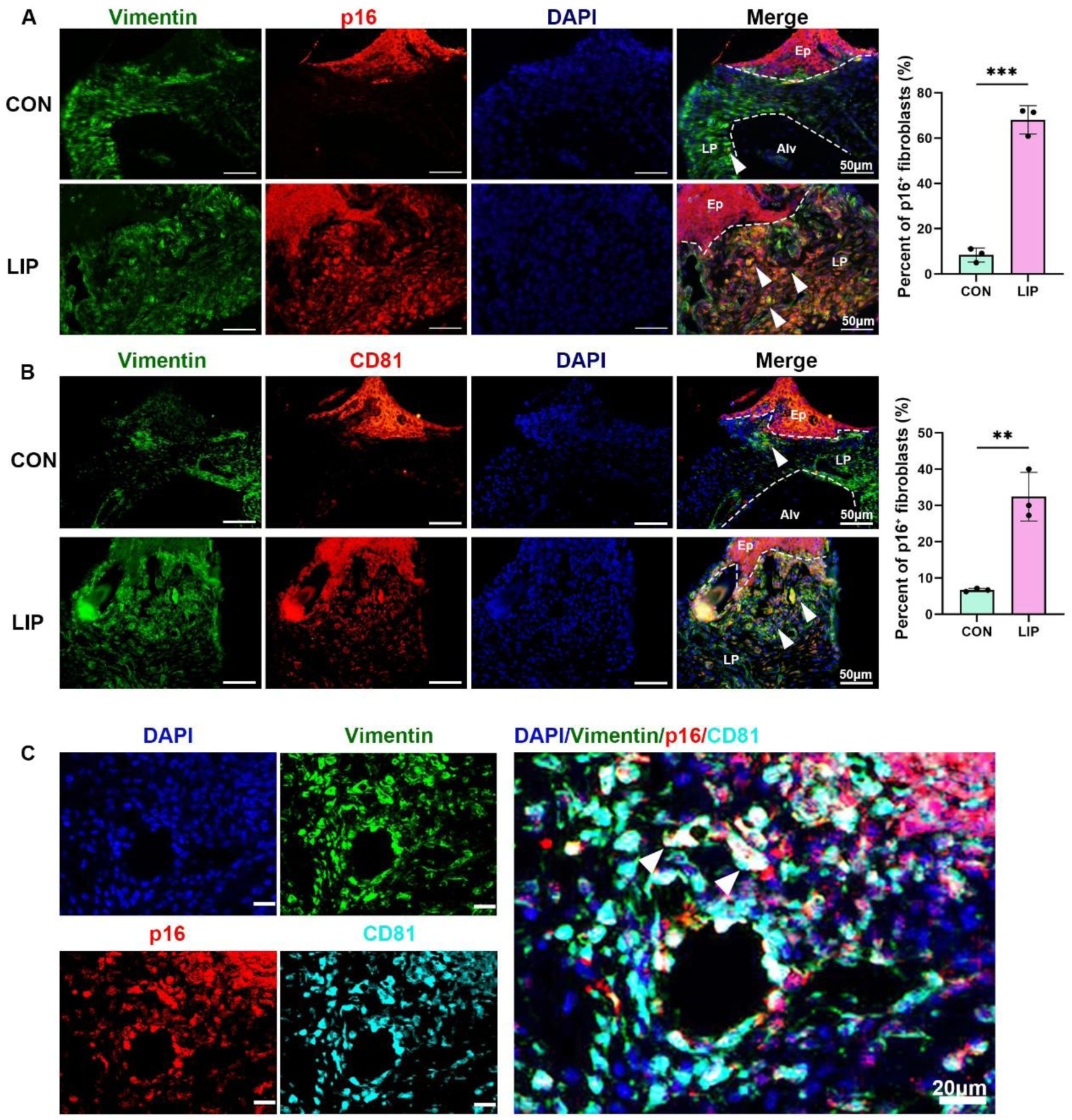
**(A)** Immunofluorescence staining and semi-quantification of p16 (red), Vimentin (green), and nuclei (blue) in control and LIP mouse gingiva, n=3 mice, scale bar=50μm. **(B)** Immunofluorescence staining and semi-quantification of CD81 (red), Vimentin (green), and nuclei (blue) in control and LIP mouse gingiva , n=3 mice, scale bar=50μm. **(C)** Immunofluorescence staining of p16 (red), VIM (green), CD81 (cyan), and nuclei (blue) in LIP mouse gingiva, scale bar=20μm. white arrow indicates triple positive cells. Ep: Epithelium; LP: Lamina propria; Alv: Alveolar bone; Data are expressed as mean ± SD. *p<0.05, **p<0.01, ***p<0.001. ****p<0.0001.

**Figure 5-figure supplement 2.**
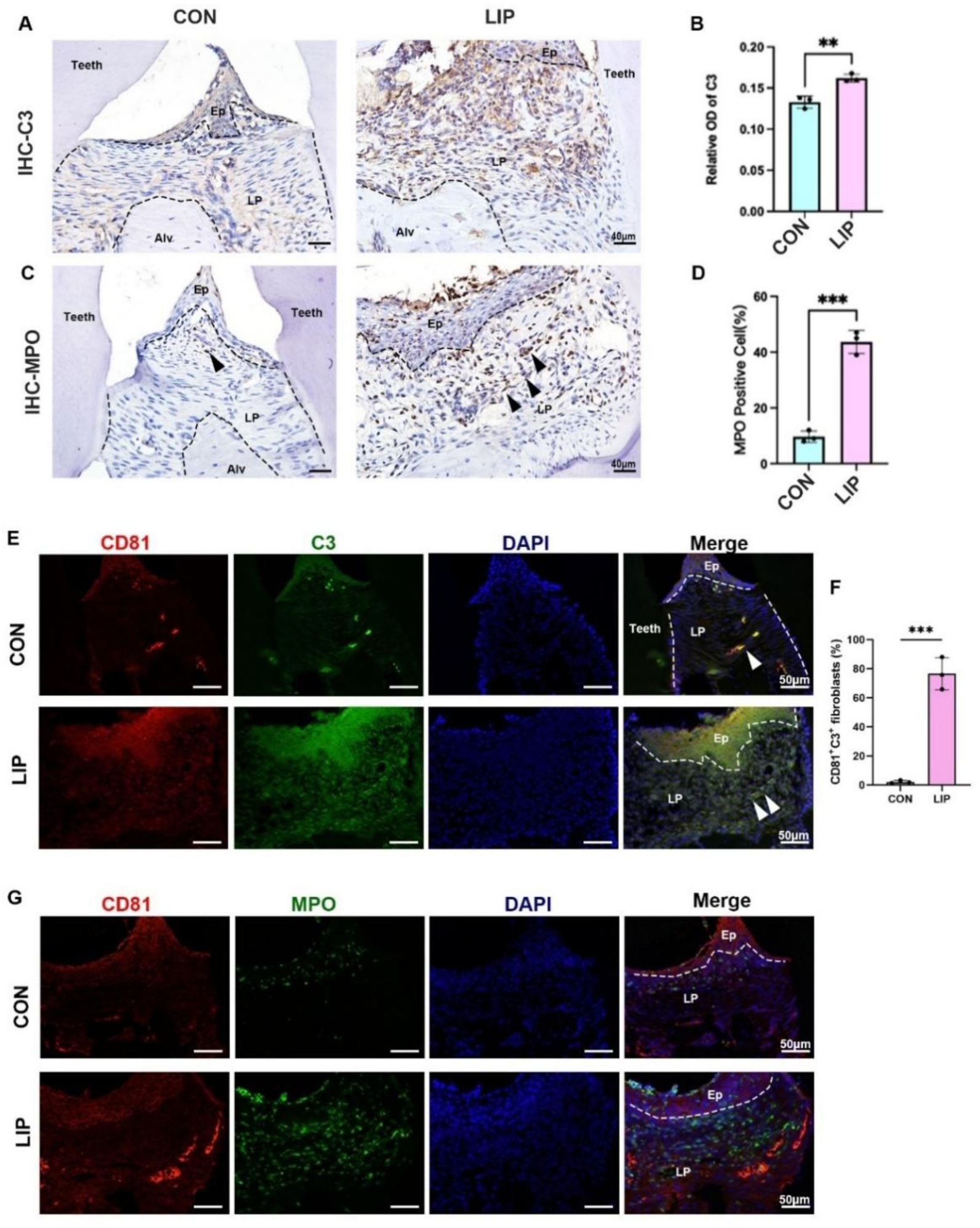
**(A)** Representative image of and **(B)** semi-quantification of IHC staining regarding C3 in control and LIP mouse gingiva. Scale bar=40μm, n=3. **(C)** Representative image of and **(D)** semi-quantification of IHC staining regarding MPO in control and LIP mouse gingiva. Scale bar=40μm, n=3. Black arrow indicates neutrophil cells. **(E)** Immunofluorescence staining and **(F)** semi-quantification of CD81 (red), C3 (green), and nuclei (blue) in control and LIP mouse gingiva, n=3 mice, scale bar=50μm. White arrow indicates double positive cells. **(G)** Immunofluorescence staining of CD81 (red), MPO (green), and nuclei (blue) in control and LIP mouse gingiva. Scale bar=50μm. Ep: Epithelium; LP: Lamina propria; Alv: Alveolar bone; Data are expressed as mean ± SD. *p<0.05, **p<0.01, ***p<0.001. ****p<0.0001.

**Figure 7-figure supplement 1.**
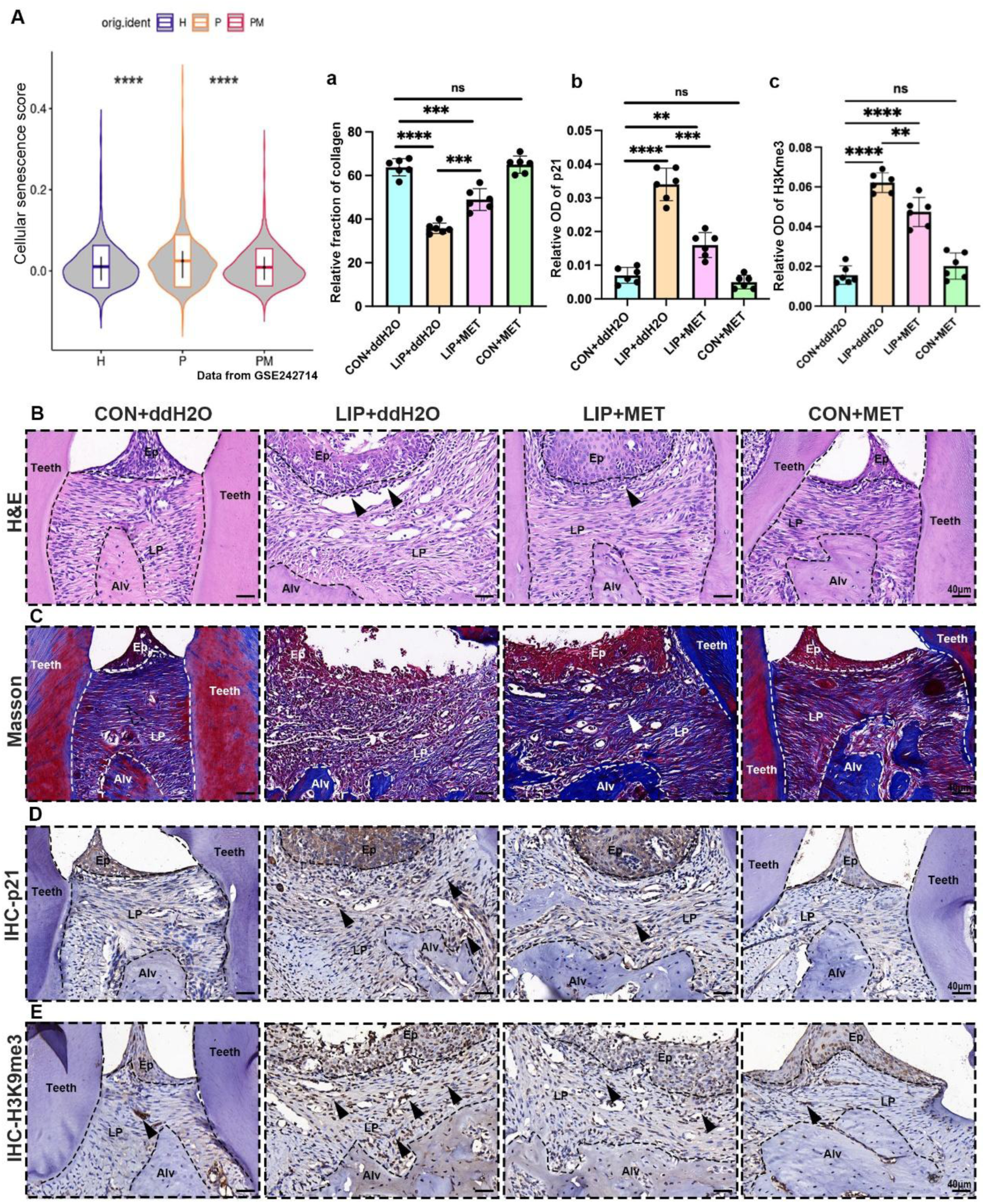
**(A)** The violin plot showing cellular senescence score in mouse gingiva of healthy (H), LIP (P) and LIP treated with metformin (PM) groups from public data GSE242714. **(B)** H&E staining images in each group. Inflammatory cells were labelled by black arrows, scale bar=40μm. **(C)** Representative image and **(a)** semi-quantification of Masson staining, in which collagen fibers were stained into blue, in each group. Collagen fiber was labelled by white arrows, n= 6, scale bar=40μm. **(D)** IHC staining and **(b)** semi-quantification of p21 in each group. Positive cells were labelled by black arrows, n=6, scale bar=40μm. **(E)** IHC staining and **(c)** semi-quantification of H3K9me3 in each group. Positive cells were labelled by black arrows, n=6, scale bar=40μm. Ep: Epithelium; LP: Lamina propria; Alv: Alveolar bone; Teeth. Data are expressed as mean ± SD. *p<0.05, **p<0.01, ***p<0.001. ****p<0.0001.

**Figure 7-figure supplement 2.**
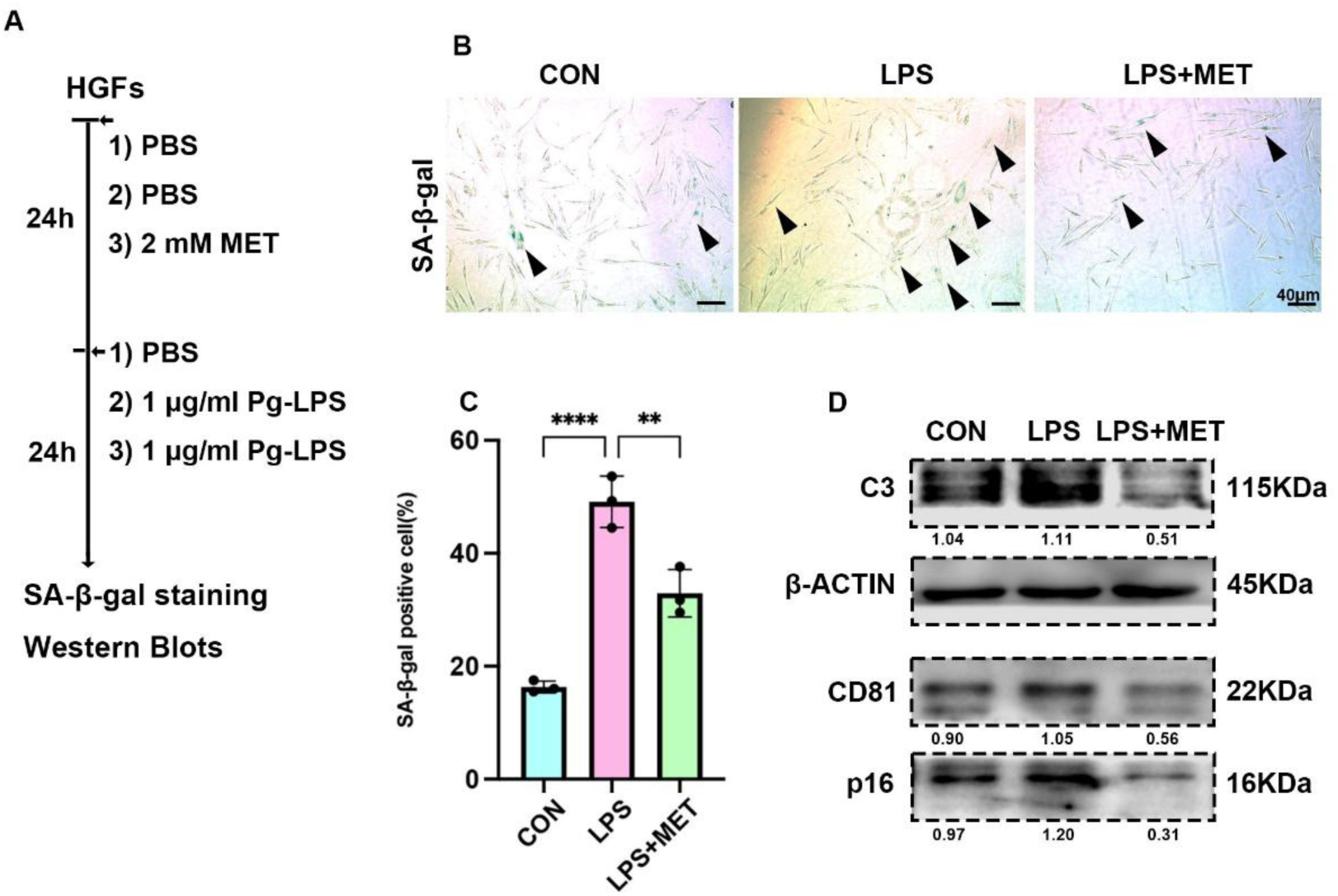
**(A)** In vitro experiment model of Pg-LPS induced senescence of human gingival fibroblasts (HGFs) treated with or without metformin (MET). **(B)** SA-β-Gal staining and **(C)** semi-quantification of Pg-LPS induced senescence of human gingival fibroblasts (HGFs) treated with or without metformin (MET). Black arrow indicates positive cells, n=3, scale bar=40μm. **(D)** Western blot image of human C3, CD81 and p16 protein levels of Pg-LPS induced senescence of human gingival fibroblasts (HGFs) treated with or without metformin (MET).

**Figure 7-figure supplement 3.**
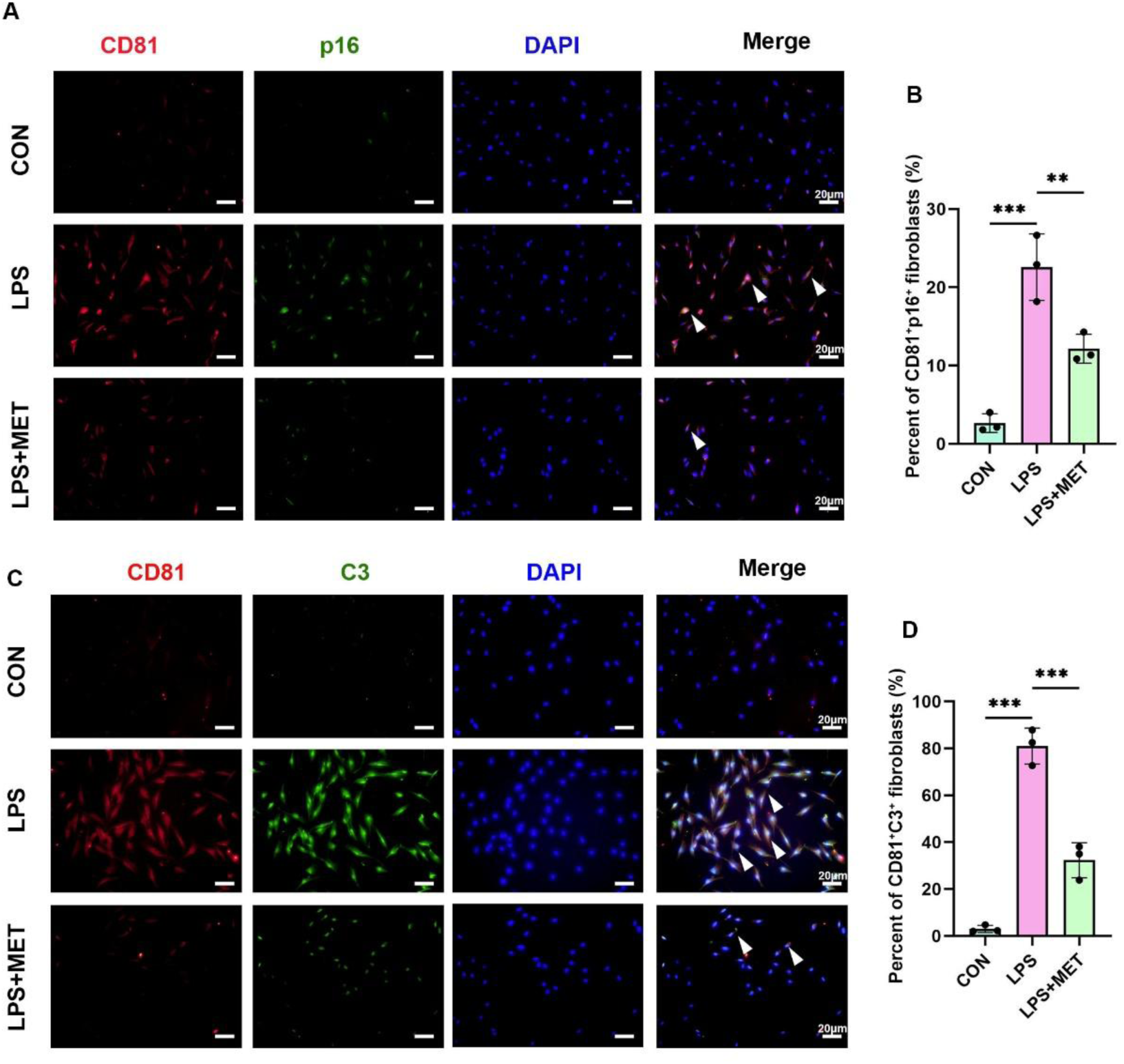
**(A)** Immunofluorescence staining and **(B)** semi-quantification of CD81^+^ p16^+^fibroblasts in Pg-LPS induced senescence treated with or without metformin (MET). CD81 (red), P16 (green) and nuclei (blue). White arrow indicates double positive cells, n=3, scale bar=20μm. **(C)** Immunofluorescence staining and **(D)** semi-quantification of CD81^+^ C3^+^fibroblasts in Pg-LPS induced senescence treated with or without metformin (MET). CD81 (red), C3 (green) and nuclei (blue). White arrow indicates double positive cells, n=3, scale bar=20μm. Data are expressed as mean ± SD.* P <= 0.05, **P<= 0.01, ***P<= 0.001, ****P<= 0.0001.

**Scheme 1.**
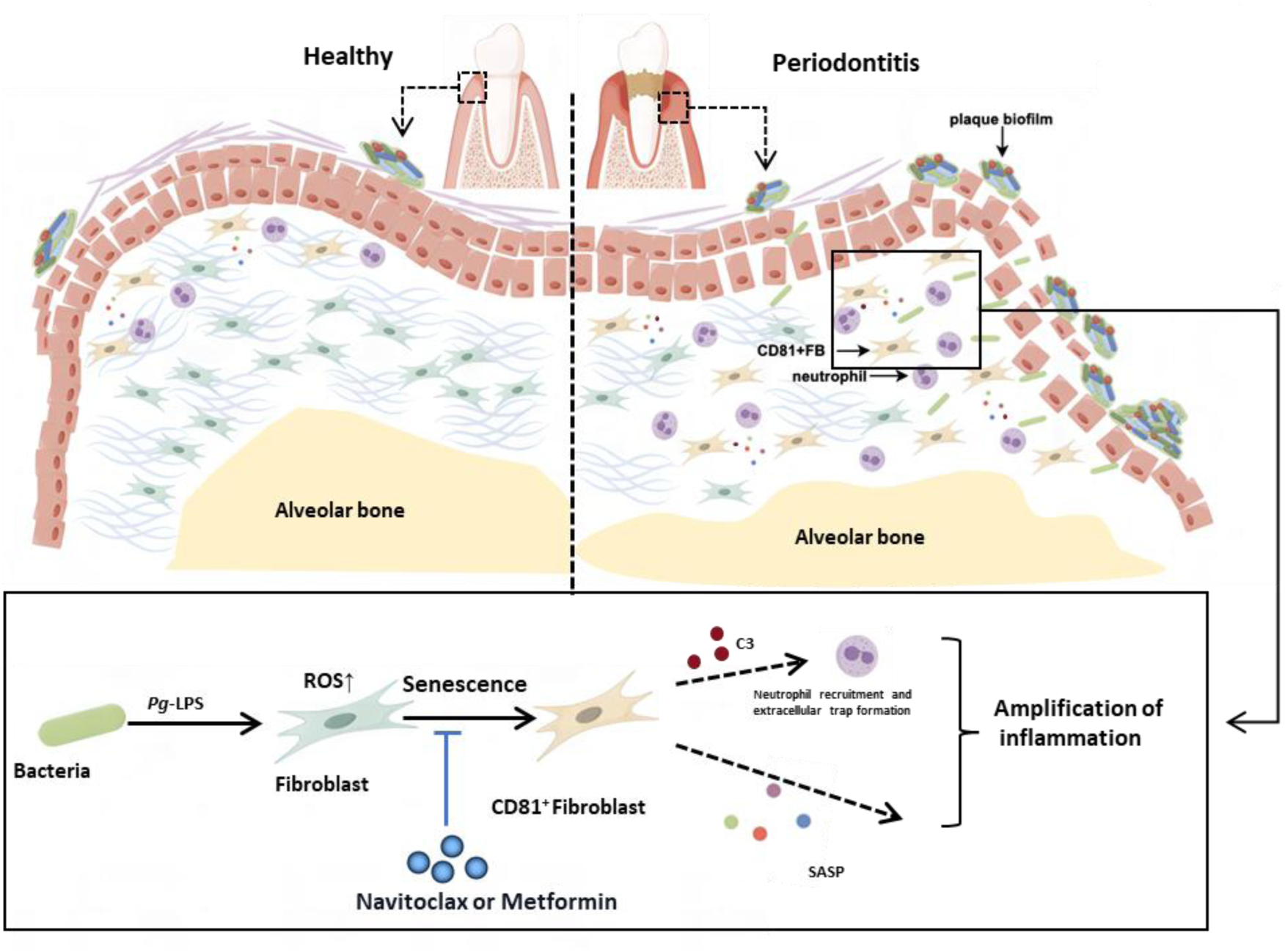
Schematic overview of the CD81^+^ senescent gingival fibroblast-neutrophil axis in periodontitis progression. We propose that the initial periodontal inflammation is triggered by the CD81^+^ senescent gingival fibroblast induced by bacterial virulence like Pg-LPS. CD81^+^ senescent gingival fibroblast could exaggerate inflammation in the periodontal tissue via secreting SASPs and recruiting neutrophils by C3. In addition, Navitoclax and Metformin could alleviate the cellular senescence of the fibroblast and rescue the uncontrolled inflammation and bone resorption.

