## Supplemental Table 1 and 2 for "CD81^+^ senescent-like fibroblasts exaggerate inflammation and activate neutrophils via C3/C3aR1 axis in periodontitis"

Post address: Department of Oral Implantology, School and Hospital of Stomatology, Wuhan University, 237 Luoyu Road, Wuhan, 430079, Hubei Province, PR China

**Declaration of Competing Interest**

The authors declare that they have no known competing financial interests or personal relationships that could have appeared to influence the work reported in this paper.

**Supplementary Table S1. Basic information of included clinical patients**

| Sample | Group | Gender | Age(year) | Usage |
|---|---|---|---|---|
| Patient 1 | Healthy | Male | 21 | Primary cell culture |
| Patient 2 | | Female | 20 | |
| Patient 3 | | Female | 33 | Histological analysis |
| Patient 4 | | Male | 25 | |
| Patient 5 | | Male | 27 | |
| Patient 6 | | Male | 28 | Frozen section |
| Patient 7 | | Male | 22 | |
| Patient 8 | | Male | 24 | |
| Patient 9 | Chronic Periodontitis | Female | 52 | Primary cell culture |
| Patient 10 | | Male | 62 | |
| Patient 11 | | Female | 60 | Histological analysis |
| Patient 12 | | Female | 50 | |
| Patient 13 | | Female | 44 | |
| Patient 14 | | Male | 25 | Frozen section |
| Patient 15 | | Male | 37 | |
| Patient 16 | | Female | 28 | |

**Supplementary Table 2.** Primer sequences for real-time PCR

| Gene | 5'-3' | Primer Sequences (5'- 3') |
|---|---|---|
| Mouse-*p16* | Forward Primer | TGTTGAGGCTAGAGAGGATCTTG |
| | Reverse Primer | CGAATCTGCACCGTAGTTGAGC |
| Mouse-*p21* | Forward Primer | TCGCTGTCTTGCACTCTGGTGT |
| | Reverse Primer | CCAATCTGCGCTTGGAGTGATAG |
| Mouse-*Tp53* | Forward Primer | GTCACAGCACATGACGGAGG |
| | Reverse Primer | TCTTCCAGATGCTCGGGATAC |
| Mouse-*β-actin* | Forward Primer | AGATGACCCAGATCATGTTTGAGA |
| | Reverse Primer | AGAGCCACCAATCCACACAG |
